# Lysosomal proteomics reveals mechanisms of neuronal apoE4-associated lysosomal dysfunction

**DOI:** 10.1101/2023.10.02.560519

**Authors:** Einar K. Krogsaeter, Justin McKetney, Leopoldo Valiente-Banuet, Angelica Marquez, Alexandra Willis, Zeynep Cakir, Erica Stevenson, Gwendolyn M. Jang, Antara Rao, Emmy Li, Anton Zhou, Anjani Attili, Timothy S. Chang, Martin Kampmann, Yadong Huang, Nevan J. Krogan, Danielle L. Swaney

## Abstract

ApoE4 is the primary risk factor for Alzheimer Disease (AD). Early AD pathological events first affect the neuronal endolysosomal system, which in turn causes neuronal protein aggregation and cell death. Despite the crucial influence of lysosomes upon AD pathophysiology, and that apoE4 localizes to lysosomes, the influence of apoE4 on lysosomal function remains unexplored. We find that expression of apoE4 in neuronal cell lines results in lysosomal alkalinization and impaired lysosomal function. To identify driving factors for these defects, we performed quantitative lysosomal proteome profiling. This revealed that apoE4 expression results in differential regulation of numerous lysosomal proteins, correlating with apoE allele status and disease severity in AD brains. In particular, apoE4 expression results in the depletion of lysosomal Lgals3bp and the accumulation of lysosomal Tmed5. We additionally validated that these lysosomal protein changes can be targeted to modulate lysosomal function. Taken together, this work thereby reveals that apoE4 causes widespread lysosomal defects through remodeling the lysosomal proteome, with the lysosomal Tmed5 accumulation and Lgals3bp depletion manifesting as lysosomal alkalinization in apoE4 neurons.

## Introduction

Alzheimer Disease (AD) is the most common form of dementia and is expected to remain a continuing major global health burden, with cases predicted to double to more than 130 million worldwide by 2050 [1]. Although the molecular and cellular complexities of AD present significant challenges for tackling this devastating disease, the apolipoprotein (apo) E4 genetic variant stands out as a predominant feature among AD cases [2, 3]. This variant, marked by the single amino acid substitution C130R, is not only found in 60-70% percent of all sporadic and familial AD cases, but is also associated with a lower age of AD onset [4–6]. Structurally, apoE4 favors intramolecular domain interaction between its N-terminal 4-helix bundle/receptor-binding domain and the C-terminal lipid binding region, taking on a closed conformation. The closed apoE4 variant thus shifts its preference from small, phospholipid-rich HDL particles to large triglyceride-rich VLDL particles [7]. ApoE4 also modifies the disease trajectory of autosomal dominant, early-onset AD, accelerating cognitive decline in *PSEN1* E280A carriers [8]. Importantly, APOE4 carriers frequently present with two hallmark pathologies of AD: Tau neurofibrillary tangles (NFTs) and β-amyloid plaques.

Considering that APOE4 is both a strong genetic risk factor for AD and is relatively common among the general population (20-25%), significant efforts have been made to understand the mechanisms by which APOE4 drives AD biology. Specifically neuronal apoE4 appears central in AD pathogenesis: While neurons typically express a minority (∼20%) of the total brain apoE [9], stress and neuronal injury results in alternative apoE splicing, facilitating its translation and increasing neuronal apoE protein production [10]. Importantly, this neuronal apoE appears central to AD pathogenesis, as its cell-type specific removal prevents both AD-associated cognitive decline [9] and hallmark AD pathology [11]. It is commonly accepted that apoE4 affects a variety of neuronal functions, reducing synaptic plasticity and density [12–15], decreasing the hippocampal volume [11, 16], and impairing learning and memory [17–21]. Importantly, these effects can be cell-type specific, with evidence for both neuronal [11, 22–25] and glial [21, 26, 27] apoE4 expression individually influencing AD related processes. At a cellular level, apoE4 interferes with cytoskeletal and microtubule structure [28–31], β-amyloid clearance [32–34], tau dynamics [11], metabolism [35–37], and endolysosomal function [15, 26, 38–41]. Several of these outcomes could be, in part, attributed to endolysosomal defects, as lysosomes clear defective organelles and larger protein aggregates.

Conversely, loss of lysosomal proteins can result in lysosomal storage and cause AD [42–44] or similar neurodegenerative diseases [45–49]. Inhibition of lysosomal function also directly leads to development of AD pathological features such as β-amyloid accumulation [50–52]. While apoE4 has previously been shown to impair endosomal trafficking and cause the accumulation of aggregate-prone proteins such as β-amyloid [53], its influence on lysosomal activity has not been explored. Besides APOE, mutations within other genetic risk factors for AD, such as rare presenilin-1 (PSEN1) mutations associated with early-onset AD, cause lysosomal mislocalization of the γ-secretase complex, resulting in an intracellular pool of aggregation-prone Aβ42 [54].

While PSEN1 is known to directly regulate lysosomal acidification and function [55], the effect of apoE4 on lysosomal homeostasis remains incompletely explored. A recent article described how exogenously administered apoE4 preferentially traffics to late endosomes/lysosomes and impairs autophagic flux [56]. Furthermore, lysosomal alkalinization has been observed in apoE4-expressing astrocytes, although its etiology and ramifications were not studied in depth [26]. To address this knowledge gap, we hypothesized that apoE4 expression intrinsically influences lysosomal function in neurons. To investigate this, we employed apoE-expressing neuronal cell lines and iPSC-derived neurons to evaluate how apoE4 expression affects lysosomal function and discover alterations to lysosomal proteins that drive apoE4-associated lysosomal defects.

## Results

### ApoE4 expression dramatically abrogates lysosomal function

While it has previously been shown that apoE4 expression interferes with astrocytic autophagy [57] and lysosomal pH [26], the effects of apoE4 expression on lysosomal function in neuronal cells, as well as the consequences of such lysosomal alkalinization, are unexplored. We set out to investigate the lysosomal influence of human apoE (hAPOE) in Neuro-2a cells, a tractable model system for studying apoE4-associated neuropathology. Importantly, physiologically relevant expression levels of hAPOE (both E3 and E4 variant) were achieved by introduction of human APOE locus flanked by 5kb and 8kb of its 5’ and 3’ genomic regions, respectively [58]. Having verified that the Neuro-2a cells expressed similar levels of apoE on a mRNA (Fig. S1A) and protein level (previous article, see [31]), we next assessed whether apoE4 influences lysosomal acidification by labeling lysosomes in hAPOE3- and hAPOE4-expressing cells using Lysotracker. The intensity of Lysotracker staining correlates with both the number of endolysosomal vesicles and their luminal acidity. We found that the Lysotracker intensity was significantly reduced in hAPOE4-expressing cells (Fig. 1A-B), suggesting either endolysosomal alkalinization or a reduction in endolysosomal density. We therefore stained the cells for the lysosomal membrane marker LAMP1, finding that the lysosomal density was not decreased in hAPOE4 cells (Fig. S1B-C). These findings suggested that defective acidification may underly our observations.

**Figure 1:**
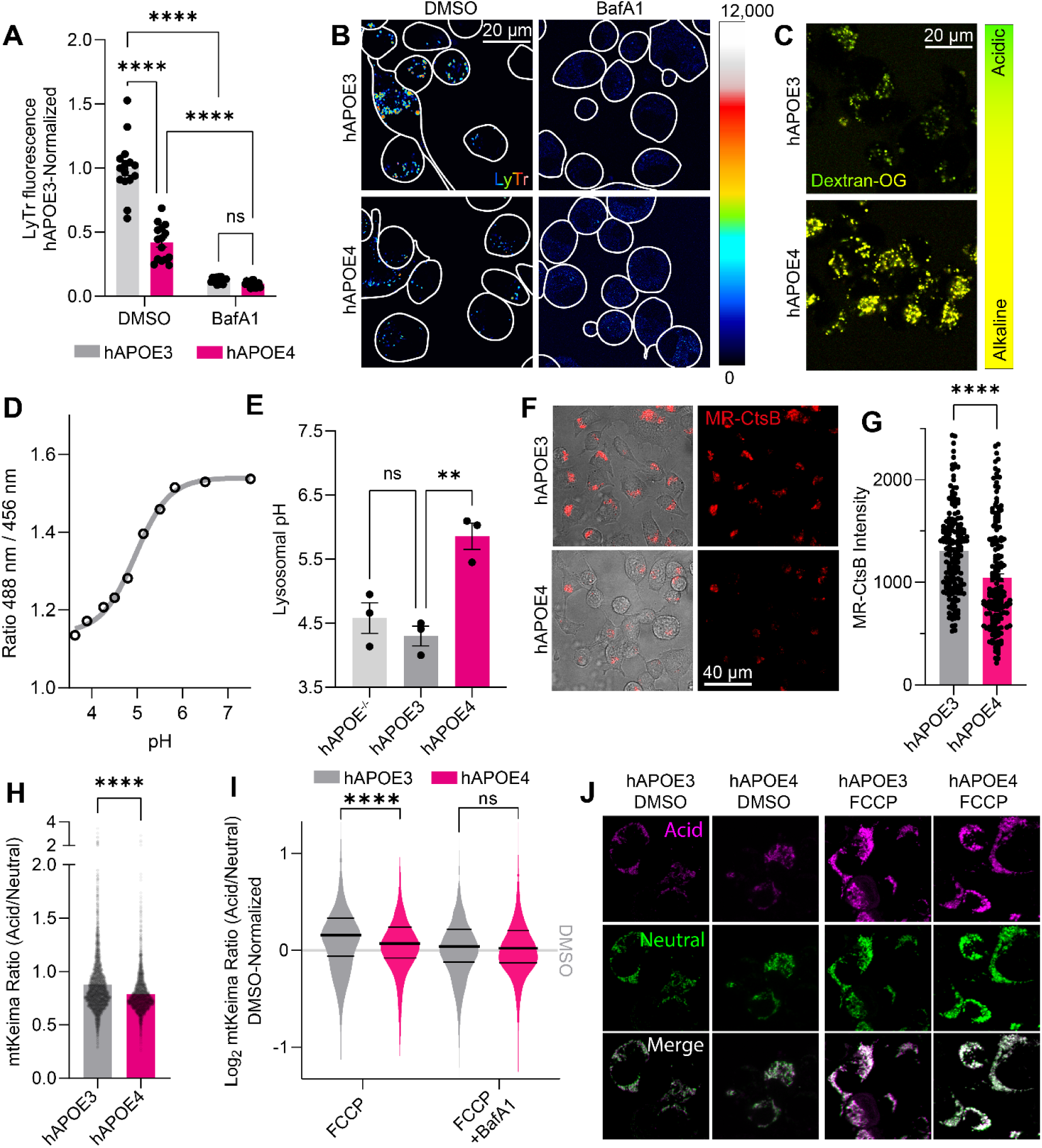
ApoE4 expression impairs lysosomal function. (**A**) LysoTracker Deep Red was used to stain acidic compartments of hAPOE-expressing Neuro-2a cells. Staining intensity was calculated per cell, and summarized for each image, across three independent experiments. Bafilomycin A1 was used to block lysosomal acidification, decreasing the lysotracker staining intensity. (**B**) Representative images for (A). (**C**) Images of lysosomal pH measurements in hAPOE-expressing Neuro-2a cells using OregonGreen-70,000 kDa Dextran. (**D**) Standard curve for the OregonGreen ratio to pH conversion. (**E**) Lysosomal pH values measured by OregonGreen experiments. Data points represent individual experiments. (**F**) Images of Magic Red-Cathepsin B (MR-CtsB)-stained hAPOE-expressing Neuro-2a cells, indicating lysosomal proteolysis. (**G**) MR-CtsB quantification of proteolytic cleavage, as shown in panel (F). Data points indicate individual mitochondria across three independent experiments. (**H**) Basal mtKeima measurements from hAPOE-expressing Neuro-2a cells. Data points represent individual mitochondria imaged across three independent experiments. (**I**) Log2-transformed mtKeima ratios normalized to genotype DMSO controls, following 1-hour mitophagic induction by FCCP and BafA1 to inhibit lysosomal acidification, plotted as a density plot of analyzed mitochondria across three independent experiments. Thick central line indicates the experimental mean, thinner flanking lines indicate the 1^st^ and 3^rd^ quartiles. (**J**) Representative images for (I). All experiments were performed as three independent technical and biological replicates, and analyzed in Fiji. The data was plotted as means with SEM error bars, and analyzed in GraphPad Prism by 2-way ANOVA followed *post-hoc* by Bonferroni’s multiple comparisons test (A,E,G,I) or an unpaired, two-tailed t-test (H).**P<0.01, ****P<0.0001.

In order to accurately measure the lysosomal pH, we employed the OregonGreen-dextran probe to ratiometrically measure the lysosomal pH. OregonGreen-dextran is routed to lysosomes by an overnight pulse, followed by a chase period allowing its endosomal (but not lysosomal) efflux. We next pH-clamped the lysosomes using a range of standard buffers to generate a standard curve of OregonGreen intensities. We could thereby extrapolate the OregonGreen intensity ratios to absolute pH values. We found that hAPOE4-expressing cells exhibit dramatic lysosomal alkalinization (pH=5.86) compared to hAPOE3 (pH=4.30) cells (Fig. 1C-E), reminiscent of previous results in apoE4-expressing astrocyte lysosomes (pH=5.20, compared to pH=4.08 in apoE3 astrocytes) [26]. This alkalinization was also observed in comparison to the parental cell line in which no human APOE had been introduced (and hAPOE^−/−^ (pH=4.58). In contrast, the lysosomal pH was comparable between hAPOE3 and hAPOE^−/−^ cells (Fig. 1E). These results suggest that hAPOE4 uniquely confers a toxic gain-of-function to lysosomes.

Maintenance of lysosomal pH is required for autophagy. We therefore assessed colocalization of lysosomes with autophagosomal LC3, and measured the cellular LC3-II/LC3-I ratios as an indicator for autophagic induction and flux. Autophagy appeared generally functional in these cells, despite the basal autophagy being slightly dampened in hAPOE4 cells (Fig. S1D-F). Since the lysosomal pH gradient is in part maintained by H^+^/Ca^2+^ antiporters [59], we also examined the lysosomal Ca^2+^ content. We targeted a previously described [60] genetically encoded calcium indicator to the lysosomes to measure lysosomal calcium release. Using GPN to release lysosomal calcium [61], we found lysosomal calcium levels to be comparable between hAPOE3 and hAPOE4-expressing cells (Fig. S1G-L), particularly after correcting for lysosomal density (Fig. S1M). These findings suggest that the apoE4-associated lysosomal alkalinization is distinct from lysosomal Ca^2+^ storage. Lysosomes normally maintain their pH at a range between 4.00 and 5.00 to support their resident, H^+^-dependent proteases. Lysosomal alkalinization naturally occurs upon ageing and cellular senescence, resulting in impaired lysosomal proteolysis and accumulation of protein aggregates. Hypothesizing that the apoE4-associated lysosomal alkalinization also impairs lysosomal degradation, we loaded the cells with Magic Red Cathepsin B (MR-CtsB), a dye that is sequestered in lysosomes and fluoresces upon its cathepsin-dependent proteolytic cleavage. We found that the hAPOE4-expressing cells exhibited significantly decreased MR-CtsB proteolysis compared to hAPOE3-expressing cells, in line with their decreased lysosomal acidity (Fig. 1F-G). Similarly, lysosomal function is crucial for the clearance of defective organelles, including mitochondrial degradation through mitophagy [60]. Mitophagy can be readily measured using the mtKeima probe [62]. Under basal conditions, we observed a modest impairment of mitophagy in hAPOE4 cells (Fig. 1H). While FCCP triggered mitophagy in both hAPOE3- and hAPOE4-expressing cells, its induction was stronger in the hAPOE3-expressing cells (Fig. 1I-J). To validate that we detected the uptake of mitochondria into lysosomes, we combined FCCP treatment with bafilomycin A1 (BafA1) to block lysosomal acidification. While BafA1 as expected reduced the acidic mtKeima signal, it intriguingly also abrogated the difference between hAPOE3- and hAPOE4-expressing cells (Fig. 1I), suggesting that mitophagic defects could be attributed to defective lysosomal acidification. Having established that neuronal apoE4 expression leads to widespread lysosomal impairments, we set out to explore the molecular mechanisms underlying this phenomenon.

### Identification of lysosomal proteins via LysoIP proteomics

Since proteins serve as principal biological effectors that many be intrinsically tied to the observed lysosomal impairment, we examined changes to the lysosomal proteome upon apoE4 expression. We used lentiviral transduction to introduce the lysosomal tag TMEM192-3xHA into each of our Neuro-2a cell lines (hAPOE^−/−^, hAPOE3, and hAPOE4) [63]. Lysosomes were visualized in transduced cells by immunostaining for the 3xHA tag, with no immunoreactivity in background cells (Fig. 2A). We immunoprecipitated the HA-tagged lysosomes (LysoIP) to isolate the lysosomal proteome, using untagged cells as a background proteome to control for non-specific binding to the anti-HA beads (Fig. 2B). Additionally, to provide greater context for alterations observed in the lysosomal proteome, we also measured protein abundances from whole cell lysates for each of the apoE genotypes. Although all cells expressed comparable levels of TMEM192, with the protein being particularly enriched in lysosomal fractions, slightly more TMEM192 was detected in hAPOE3 lysosomes (Fig. S2A). Overall, the lysosomal protein entities detected, and the relative frequencies of these, were comparable across genotypes (Fig. S2B-C). As expected, clustering of protein abundances across all samples revealed distinct proteome signatures between the three different experimental groups (whole cell lysate, LysoIP, and background), with no discernible influence of overexpressing the lysosomal TMEM192-3xHA tag on the whole-cell proteome (Fig. 2C; full heat-maps shown in Fig. S3A-B; Table S1A). Additionally, compared to the background samples, our LysoIP samples were appropriately enriched in proteins with lysosomal annotations as compared to other subcellular organelles (Fig. 2D). Next, we focused on defining the lysosomal proteome. First, we identified enriched lysosomal proteins by comparing the tagged and background samples for each genotype. We considered proteins to be lysosomal if they were either exclusively detected in LysoIP samples, or significantly enriched in LysoIP samples relative to background samples (>2-fold, and p<0.05) (Fig. 2E; Table S1B-C). Gene set enrichment analysis of these proteins highlighted an enrichment of lysosome-associated terms, including autophagy, endosome membrane, late endosome, and vacuolar membrane (Fig. 2F). As expected, the most significantly enriched proteins included well-known lysosomal proteins, including degradative enzymes (Ctsd, Psma3), proton pump subunits (Atp6v1e1), metabolic enzymes (Gusb), and lysosomal scaffold proteins (Lamtor 2/4). However, we also found a number of proteins to be detected exclusively and reproducibly across all lysosomal samples, but that have not yet been ascribed a lysosomal function. These include constituents of the Usp8:Ptpn23:Stam2 and Nedd4:Litaf:Bmpr1 deubiquitin- and ubiquitination complexes, and the early endosome/Rab5a-associated proteins Rabep1 and Rabgef1. While neither complex is appreciated to function in the lysosome, their respective implicated processes of protein ubiquitination and Rab protein activation could conceivably be of lysosomal importance. Lastly, the primary driver of differences between LysoIP samples, as determined by principal component analysis (PCA), was found to be bafilomycin treatment, (Fig. S4A), and was distinguished by lysosomal accumulation of chaperonin TCP-1 subunits, proteasome subunits, and the dynein complex, alongside loss of cathepsins, the GATOR2 complex, and glycosyl hydrolases (Fig. S4B; Table S1C).

**Figure 2:**
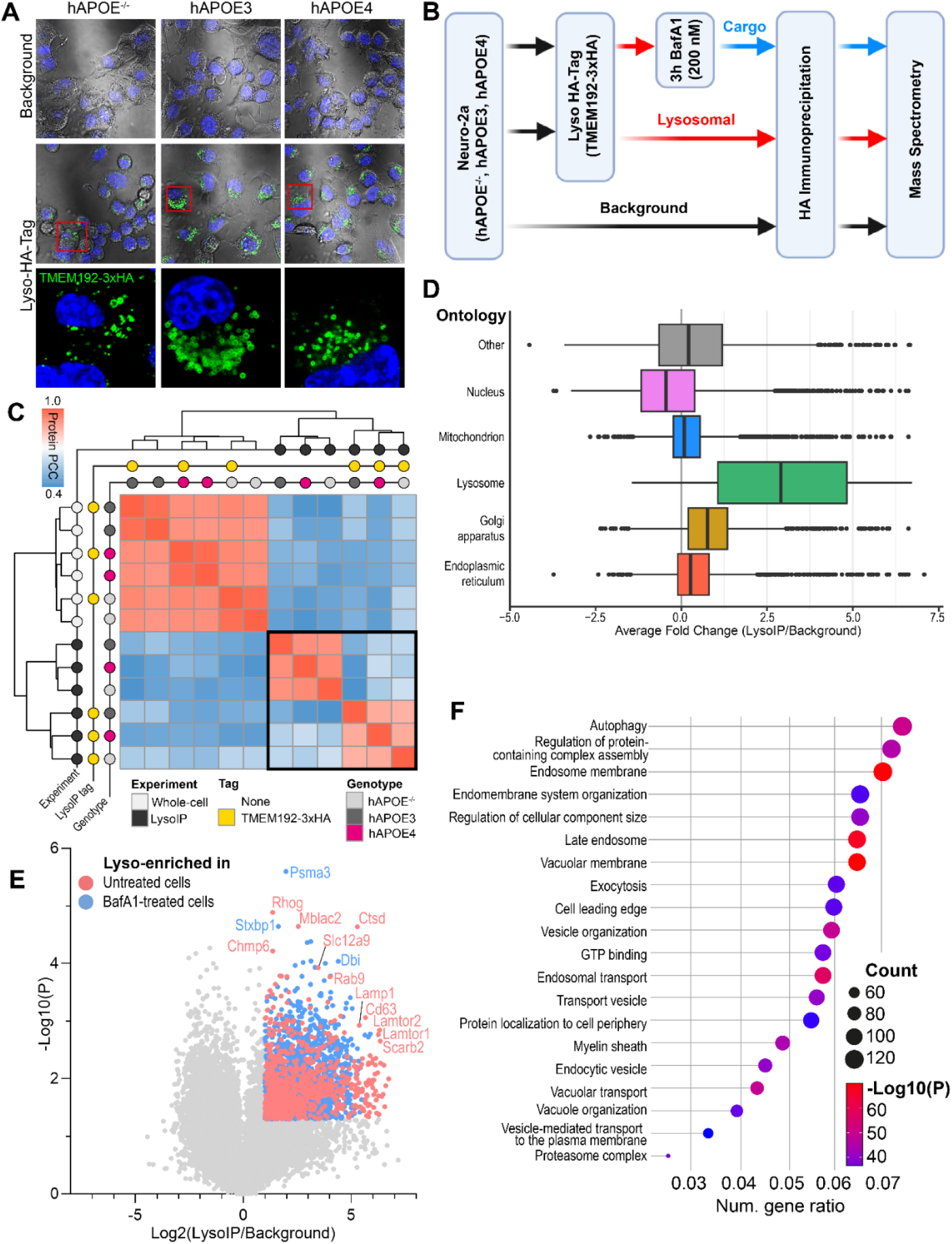
Isolation of lysosomes by lysosomal immunoprecipitation (LysoIP) (**A**) Immunostaining of the lysosomal TMEM192-3xHA tag shows immunoreactivity only in polyclonal, transduced Neuro-2a cells. (**B**) Schematic of the LysoIP workflow. Note, paired samples from whole cell lysates of each of these conditions were also analyzed. (**C**) Clustered heatmap of protein similarities (measured by global protein Pearson’s correlation coefficient; PCC) between sample sets reveals clustering of whole-cell, LysoIP background and LysoIP-tagged samples. Sample sets within the black bounding box are further analyzed in panels (D-E). (**D**) Box plot of protein abundances in the untreated LysoIP samples relative to background IP samples. Proteins have been separated based on their subcellular compartment ontologies. (**E**) Volcano plot of untreated or Baf1A treated LysoIP samples as compared to background IP samples. Colored dots indicate proteins with the minimum enrichment (Log2(LysoIP/Background)>1 and p<0.05) to be categorized as lysosomal proteins. (**F**) Gene ontology enrichment of Neuro-2a lysosomal proteins (either exclusively detected in LysoIP samples or significantly enriched in LysoIP samples relative to background, as indicated in panel E). Dot size indicates the number of lysosomal proteins matching to that ontology term, while the color represents the enrichment of that ontology term.

### Proteomic alterations to apoE4-associated lysosomes

After demonstrating successful enrichment of lysosomal proteins, we next investigated the consequences of apoE4 expression on the lysosomal proteome, particularly in light of the lysosomal disturbances observed in hAPOE4-expressing cells (Table S1D-E). Lysosomal proteomes could be readily separated by genotype both by principal component analysis (PCA; Fig. 3A) and hierarchical clustering (Fig. S3A-B; Fig. 2C). PCA component 2 (11.6% variance) separated hAPOE3- and hAPOE4-expressing samples based on hAPOE4-associated accumulation of glucosyl hydrolases, Lamtor proteins, and a number of Golgi-associated vesicle proteins, while enzymes regulating aminoacyl-tRNA biosynthesis and gluconeogenesis were depleted. Component 3 (7.4% variance) separated hAPOE^−/−^ lysosomes from hAPOE-expressing lysosomes, marked by hAPOE-dependent accumulation of sterol transporters and proton pump subunits, and a depletion of proteasome subunits and microtubule proteins (Fig. 3A). Comparing hAPOE4- to hAPOE3-expressing cells, we observed an increase in TMED proteins (Fig. 3B), implicated in ER-Golgi trafficking and sorting misfolded proteins to the lysosomes [64]. Intriguingly, most lysosomal proteins with lower abundance in hAPOE4 cells were Bafilomycin A1-sensitive proteins that accumulate upon proton pump blockade, including the phospho-Tau-binding protein Scrn1 (Fig. 3B) [65]. The few intrinsically lysosomal proteins expressed at lower levels in hAPOE4 lysosomes include the ADP ribosylation factor activator Arfgef3/Big3 regulating lysosomal trafficking [66] and the glycoprotein Lgals3bp/90K-BP/Mac-2BP (Fig. 3B).

**Figure 3:**
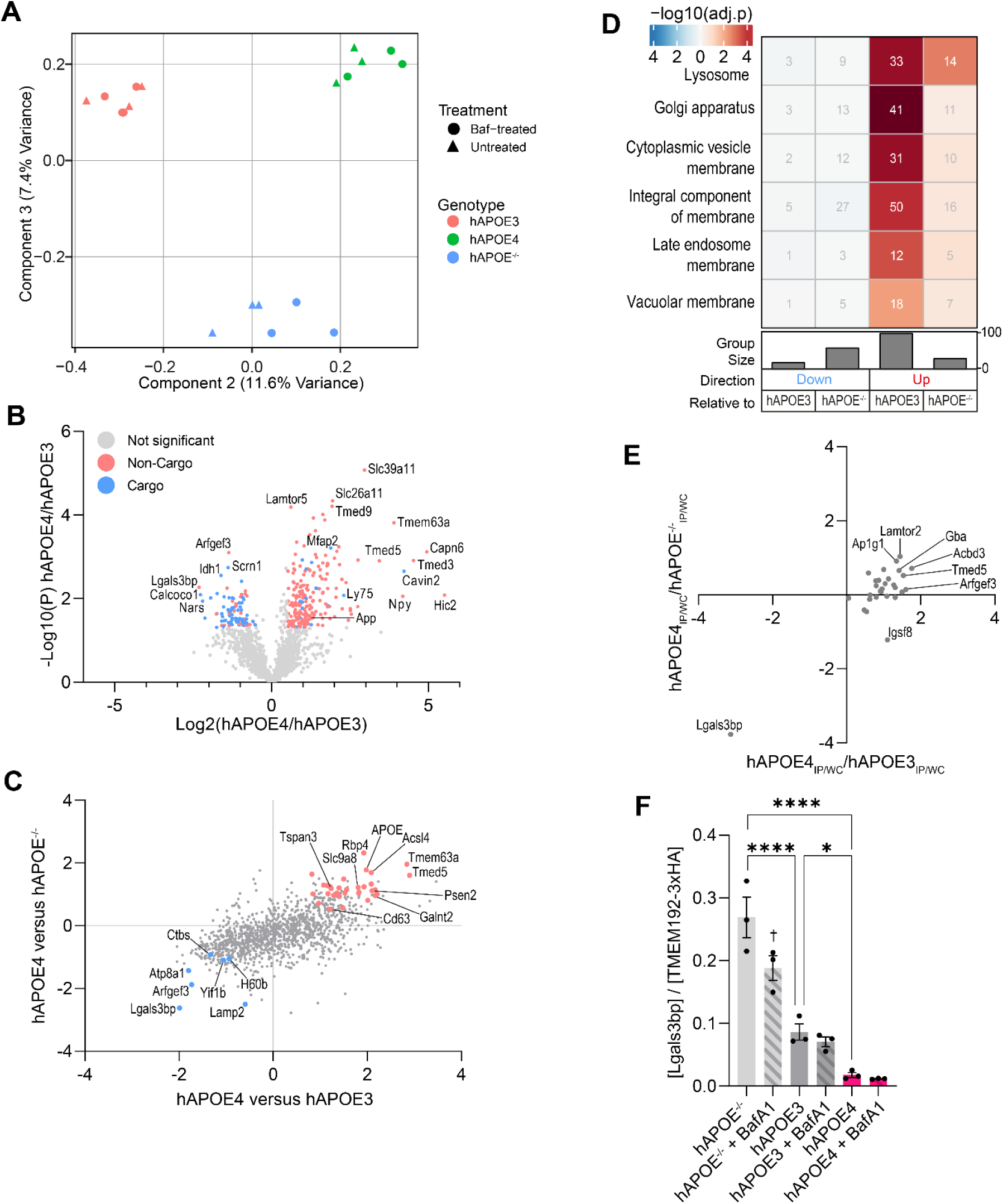
The apoE4-associated lysosomal proteome. (**A**) Principal component analysis separates Neuro-2a lysosomal proteomes by hAPOE genotype based on PCA components 2 and 3. (**B**) Volcano plot showing hAPOE4-associated lysosomal protein changes relative to hAPOE3 controls. Proteins significantly changed in hAPOE4 lysosomes are colored by their status as a lysosomal cargo protein, based on basal (red) or BafA1-dependent (blue) lysosomal enrichment. (**C**) Correlation of hAPOE4-associated lysosomal protein changes relative to both hAPOE3 and hAPOE^−/−^ controls. Axes are geometric means of Log2(hAPOE4/control) and −Log10(P) values, calculated using the formula 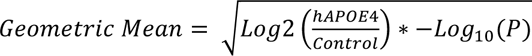. Colored proteins were statistically significantly (p < 0.05) depleted (blue) or accumulated (red) in hAPOE4 lysosomes relative to both controls. (**D**) Enrichment of gene ontology (GO) terms associated for hAPOE4-associated lysosomal proteins as compared to hAPOE3 or hAPOE^−/−^ lysosomal proteins. (**E**) Lysosomal protein distribution coefficients ([Protein]_Lysosome_/[Protein]_Whole-cell_) in hAPOE4 lysosomes relative to hAPOE3 and hAPOE^−/−^ controls. (**F**) Protein abundance of lysosomal Lgals3bp as factor of hAPOE expression, allele, and BafA1 proton pump blockade. The data was analyzed in GraphPad Prism by 2-way ANOVA followed *post-hoc* by Bonferroni’s multiple comparisons test. *P<0.05; ****P<0.0001.

Since we (and others) have found that neuronal apoE4 confers a gain-of-toxicity effect not observed in hAPOE^−/−^ or hAPOE3 cells [9, 24, 67], we extended our comparisons of the lysosomal proteome to hAPOE^−/−^ cells to identify lysosomal proteins differentially regulated uniquely in apoE4 cells. We considered proteins significant if they changed in the same direction and reached statistical significance relative to both hAPOE^−/−^ and hAPOE3 controls. To illustrate the hAPOE4-dependent proteomic alterations relative to both controls simultaneously, we calculated the geometric mean of the P and log2(hAPOE4/control) values. Proteins that reached statistical significance relative to both controls are highlighted in Fig. 3C. Through this approach, we found that hAPOE4-expressing lysosomes accumulated APOE itself, AD-associated secretase proteins (Psen2, Cd63, Tspan3), trafficking adaptors (Ap1s1, Ap1g1, Bet1l, Lamtor1, Lamtor2, Tmed5), and lipid/glycan-regulators (Acsl4, Acbd3, Galnt2, Man2b1, Neu1, Pip4p1, Sacm1l, Tmem55b, Uxs1) (Fig. 3B-C, Fig. S4C). Proteins accumulating in hAPOE4 lysosomes were principally enriched in the “Lysosome” gene ontology (GO) term, even when correcting for the already lysosome-laden background proteome (Fig. 3D). Proteins annotated under the Golgi apparatus, particularly chaperones and proteins ensuring its structural integrity, also accumulated in hAPOE4 lysosomes (Fig. 3D). Fewer changes were observed in the opposite direction, with the hAPOE4-expressing lysosomes expressing lower levels of the aforementioned Arfgef3 and Lgals3bp among others (Fig. 3B-C, Fig. S4C-D).

Since lysosomal protein changes could be secondary to whole-cell expression changes, we correlated the expression level of dysregulated lysosomal proteins to their whole-cell abundance. As expected, we observed a robust correlation between hAPOE4-associated global and lysosomal protein level changes (Fig. S4E-F; Table S1F). We hypothesized that proteins that were over- or underrepresented on hAPOE4 lysosomes could reflect changes in their lysosomal uptake and/or clearance. For example, a lysosome-accumulating protein that decreases on the whole-cell level could reflect its increased trafficking to the lysosome, resulting in lysosomal degradation. Conversely, a protein depleted from lysosomes but accumulating in the whole-cell fraction could suggest impaired lysosomal uptake, and result in its cellular accumulation. To explore such possibilities, we plotted lysosomal under- or overrepresented proteins in the context of the hAPOE4 genotype. Note, proteins not detected in whole-cell samples could not be plotted, precluding similar analyses of 15 proteins including Psen2 and Rbp4. This approach highlighted the unique case of Lgals3bp, a protein whose expression was reduced in lysosomes despite unchanged whole-cell expression levels (Fig. 3E). In contrast, several proteins were overrepresented in the lysosome, including Lamtor2, Ap1g1, Acbd3, and Tmed5, although these changes were less extreme. Performing a partial least squares discriminant analysis to identify proteins characteristically separating lysosomes of one genotype from others further highlighted Lgals3bp depletion to be a characteristic feature of hAPOE4-expressing lysosomes (Fig. S4G). Intriguingly, lysosomal Lgals3bp levels appeared sensitive to both hAPOE and proton pump blockade, with hAPOE^−/−^ lysosomes boasting the highest Lgals3bp levels and BafA1 treated hAPOE4 lysosomes containing the least Lgals3bp (Fig. 3F). This approach thereby revealed a set of hAPOE4-dysregulated lysosomal proteins where lysosomal protein abundance changes were not indirectly driven by changes in whole cell expression, but rather directly driven by changes to how these proteins are recruited to and cleared from the lysosome.

### Identification of drivers underlying lysosomal defects in apoE4 neurons

Having identified hAPOE4-sensitive lysosomal proteins, we assessed whether knockdown of these proteins affects lysosomal function. We targeted Lgals3bp as the only downregulated protein in hAPOE4 lysosomes, being the principal characteristic protein separating hAPOE4 lysosomes from hAPOE^−/−^ and hAPOE3 lysosomes. We also targeted three major AD risk factor proteins that we found to be accumulating in hAPOE4 lysosomes, specifically apoE itself, App [68] and Psen2 [52]. We also targeted tau, as it interfaces with and spreads via the autophagic/lysosomal system [69–77], and is implicated across neurological disorders (tauopathies). Finally, we targeted Tmed5, which we observed to be one of the most significantly accumulating lysosomal proteins associated with Golgi apparatus functions.

We first validated the efficacy of our siRNAs in reducing transcript levels (Fig. S5A), and protein levels of our top candidates (Fig. S5B-C). We then assessed their influence on lysosomal function. We considered an accumulating lysosomal protein a hit if its knockdown increased Lysotracker staining in hAPOE4 cells, or a depleted lysosomal protein (Lgals3bp) if its removal reduced Lysotracker staining in hAPOE3 cells. This highlighted that the apoE4-associated lysosomal impairments could be driven by Lgals3bp depletion and Tmed5 accumulation (Fig. S5D).

### ApoE4-associated lysosomal Lgals3bp depletion decreases the lysosomal density of neurons

We found that Lgals3bp depletion markedly decreased the Lysotracker staining in healthy hAPOE3 Neuro-2a cells (Fig. 4A-B). To assess whether this was driven by changes in lysosomal acidification or the density of lysosomes, we loaded the Neuro-2a cells with Dextran-Oregon Green, finding that Lgals3bp knockdown modestly alkalinized the lysosomal pH, but nearly halved the lysosomal density (Fig. 4C-E). To assess whether these changes were accompanied by altered lysosomal processing, we measured lysosomal proteolysis using MR-CtsB. In line with its modest influence on lysosomal acidification, we could not detect significant proteolytic impairments upon Lgals3bp knockdown (Fig. 4F-G). In order to examine if LGALS3BP depletion similarly decreases LysoTracker staining in post-mitotic, apoE3 human neurons, we used CRISPRi-encoding iPSCs [78, 79] to generate mature neurons (iNeu). As a positive control, we inhibited TMED10, which previously has been shown to enhance lysotracker staining [79]. We confirmed that while TMED10 inhibition increased lysotracker staining in iPSC-derived neurons, LGALS3BP inhibition decreased staining, suggesting that LGALS3BP is necessary to maintain lysosomal integrity across neuronal models (Fig. 4H-I).

**Figure 4:**
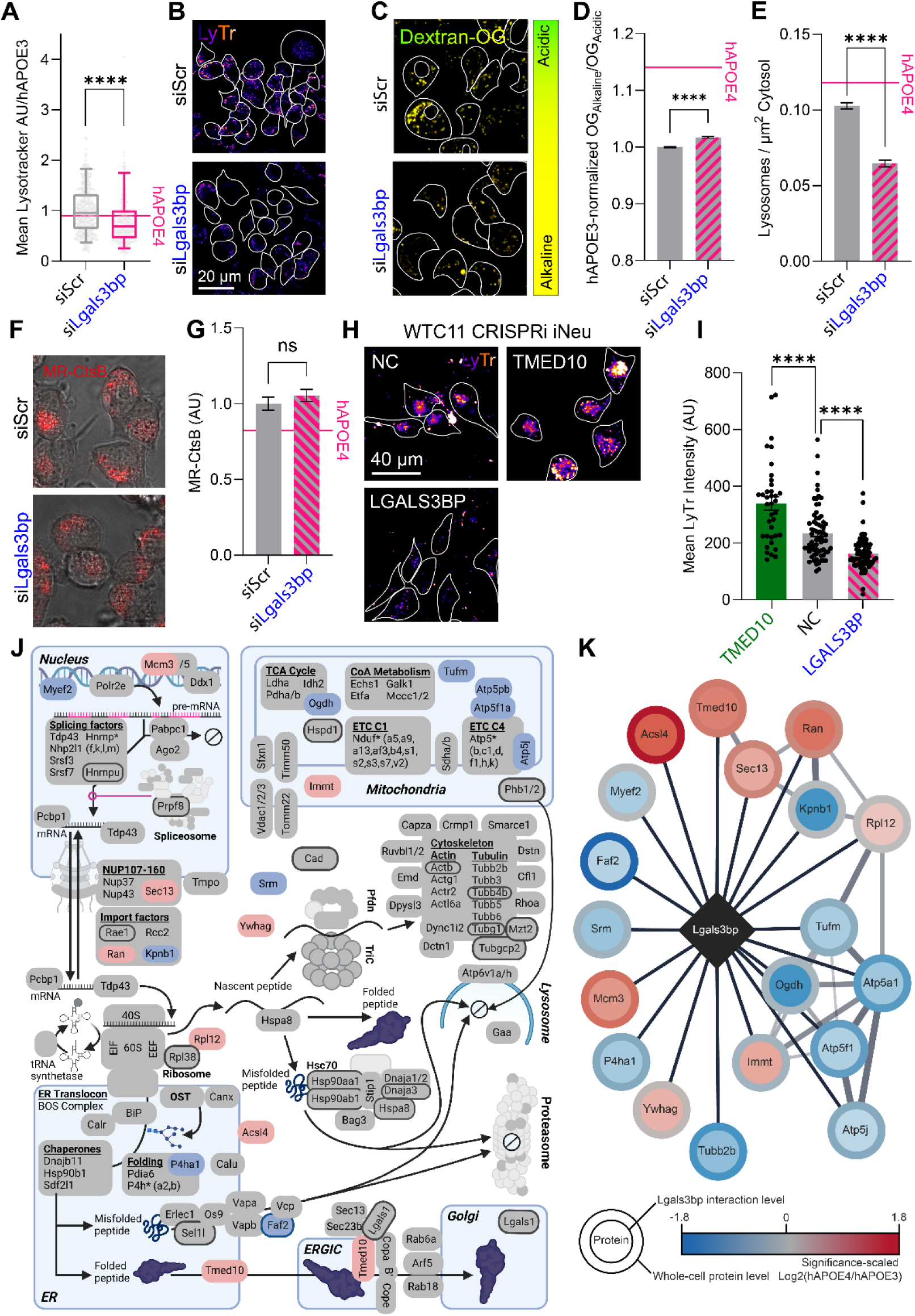
Local Lgals3bp depletion in hAPOE4 lysosomes affects the lysosomal density. (**A**) Quantification of LysoTracker staining of acidic compartments in hAPOE3 Neuro-2a cells following knockdown of Lgals3bp. Boxes indicate the 5-95 percentile. Data points indicate individual cells from three independent experiments. (**B**) Representative LysoTracker images for quantification shown in (A). (**C**) Oregon-green relative pH measurements upon Lgals3bp knockdown in hAPOE3 Neuro-2a cells, compared to scrambled controls. (**D-E**) Quantification of relative lysosomal pH (D) and density (E) upon Lgals3bp knockdown, as shown in panel (C). The value of these measurements in hAPOE4 Neuro-2a cells upon treatment with scrambled controls is shown for reference. Data is plotted as pH per lysosome or cellular lysosomal density across three independent experiments. (**F**) Magic Red Cathepsin B images of hAPOE3 Neuro-2a cells following Lgals3bp knockdown. (**G**) Quantification of Magic Red Cathepsin B intensity per cell across three independent experiments, as shown in panel (F). The value of this measurement in hAPOE4 Neuro-2a cells upon treatment with scrambled controls is shown for reference. (**H**) Representative LysoTracker images of iPSC-derived neurons following CRISPRi suppression of indicated genes, relative to untransduced negative controls (NC). (**I**) Quantification of LysoTracker measurements per cell from panel H, measured across three independent experiments. (**J**) Proteins found to bind Lgals3bp are highlighted in their canonical cellular compartment. Grey interactors were unchanged by apoE genotype; colored protein nodes indicate preferential association with Lgals3bp in the context of apoE3 (blue) or apoE4 (red) genotypes. Nodes with borders are annotated Lgals3bp interactors by the BioGRID protein interaction repository [95]. The illustration was created with BioRender.com. (**K**) Cytoscape network of apoE genotype-associated Lgals3bp interactors. Fill colors represent relative retrieval of Lgals3bp interaction partners, with red indicating interactors with greater Lgals3bp binding in hAPOE4 Neuro-2a cells, and blue indicating interactors with greater Lgals3bp binding in hAPOE3 cells. Border colors represent relative protein abundances (hAPOE4/hAPOE3) in the whole-cell lysate. All experiments were performed as three technical and biological replicates. Fluorophore intensities and spot counting was performed in Fiji. Data was analyzed in GraphPad Prism by 2-way ANOVA followed *post-hoc* by Dunnett’s multiple comparisons test. ****P<0.0001.

While Lgals3bp has been explored in the context of centriolar function and its extracellular secretion [80–83], little is known about how it affects lysosomal function. It structurally consists of three domains: An N-terminal Scavenger Receptor Cysteine-Rich (SRCR) domain associated with phagocytosis of negatively charged ligands such as lipoproteins [84], a BTB domain that recruits ubiquitin ligase complexes, and a C-terminal BTB- and Kelch-associated BACK domain [85]. This domain architecture suggests that Lgals3bp could scaffold SRCR substrates and ubiquitin ligases associated with the BTB-BACK domain, facilitating substrate ubiquitination and degradation. While Lgals3bp was recently described as a ubiquitous lysosomal protein [86], the lysosomal function of Lgals3bp is largely unknown [86].

We set out to identify the determinants of the intracellular Lgals3bp mislocalization underlying the hAPOE4-associated lysosomal depletion, and to identify putative Lgals3b-associated mediators of lysosomal function. In order to do so, we evaluated the Lgals3bp interactome by affinity-purification mass spectrometry (APMS; Table S2A-C). The obtained Lgals3bp interactomes readily clustered away from the GFP interactomes, indicating a distinct interactome not governed by the GFP affinity tag (Fig. S6D). Having identified Lgals3bp interactors, we used STRING [87] to annotate protein complexes within the Lgals3bp interaction network (Fig. S6E). Lgals3bp-associated proteins and protein complexes were next manually assigned subcellular compartments (Fig. 4J). While studies have previously focused on its functions as a secretory [81–83] or centriole-associated protein [80], Lgals3bp is associated with a diversity of protein complexes associated with the nucleus, cytoskeleton, mitochondria, the endoplasmic reticulum (ER), and lysosomes. Of particular interest, we find that Lgals3bp interacts with ER-to-lysosome-trafficking proteins, and proteins regulating lysosomal acidification (specifically Atp6v1a and Atp6v1h).

A plausible explanation for the apoE4 allele-associated Lgals3bp depletion would be a changed Lgals3bp interactome. Specifically, lower expression of its lysosome-routing proteins would result in the observed lysosomal depletion. We therefore assessed whether apoE4 expression affected the Lgals3bp interactome. Among a few altered interactors, we found that apoE4 favored interactions between Lgals3bp and the ER-to-Golgi cargo receptor Tmed10, associated with a trend towards higher Tmed10 expression (Fig. 4J-K; Table S2D-E). Conversely, the apoE4 allele context decreased the interaction of Lgals3bp with several proteins, including Faf2 which is associated with the lysosome-routing Vcp/ERAD machinery (Fig. 4J-K). Of note, Faf2 protein abundance is significantly lower in hAPOE4 cells, which may serve as a mechanism to prevent the association of Lgals3bp with this lysosome-routing interaction partner (Fig. 4K; Table S2F). Taken together, we find that several Lgals3bp interaction partners are differentially regulated in hAPOE3 and hAPOE4 cells, which has the potential to influence the localization and trafficking of Lgals3bp.

### Increased Tmed5 expression causes lysosomal overflow and alkalinization in apoE4 neurons

In contrast to Lgals3bp, we found that apoE4-associated Tmed5 accumulation in lysosomes caused lysosomal defects that could be reversed by its acute removal (Fig. 5A-B). We confirmed by lysosomal Oregon-green pH measurements that knockdown of Tmed5 significantly enhanced lysosomal acidification in hAPOE4 Neuro-2a cells, while not impacting the lysosomal density (Fig. 5C-E). We subsequently continued to assess how this influence on lysosomal acidification affected lysosomal processing. We found that Tmed5 removal significantly enhanced lysosomal proteolysis of the MR-CtsB indicator in hAPOE4 cells (Fig. 5F-G). Concurrently, we found that Tmed5 knockdown significantly enhanced mitophagy in hAPOE4 Neuro-2a cells, shifting mitochondria towards more acidic environments both under basal and FCCP-induced conditions (Fig. 5H-I). Conversely, in hAPOE3 Neuro-2a cells we found that TMED5 overexpression caused the opposite phenotype, resulting in decreased LysoTracker staining (Fig. 5J-K) and lysosomal proteolysis (Fig. 5L-M). These findings indicate an intricate link between TMED5 expression levels, where its apoE4-associated overexpression causes lysosomal alkalinization and impairing lysosomal catabolism. They also corroborate TMED5 as a tunable target for modulating lysosomal activity, with its removal alleviating lysosomal impediments observed in apoE4 cells.

**Figure 5.**
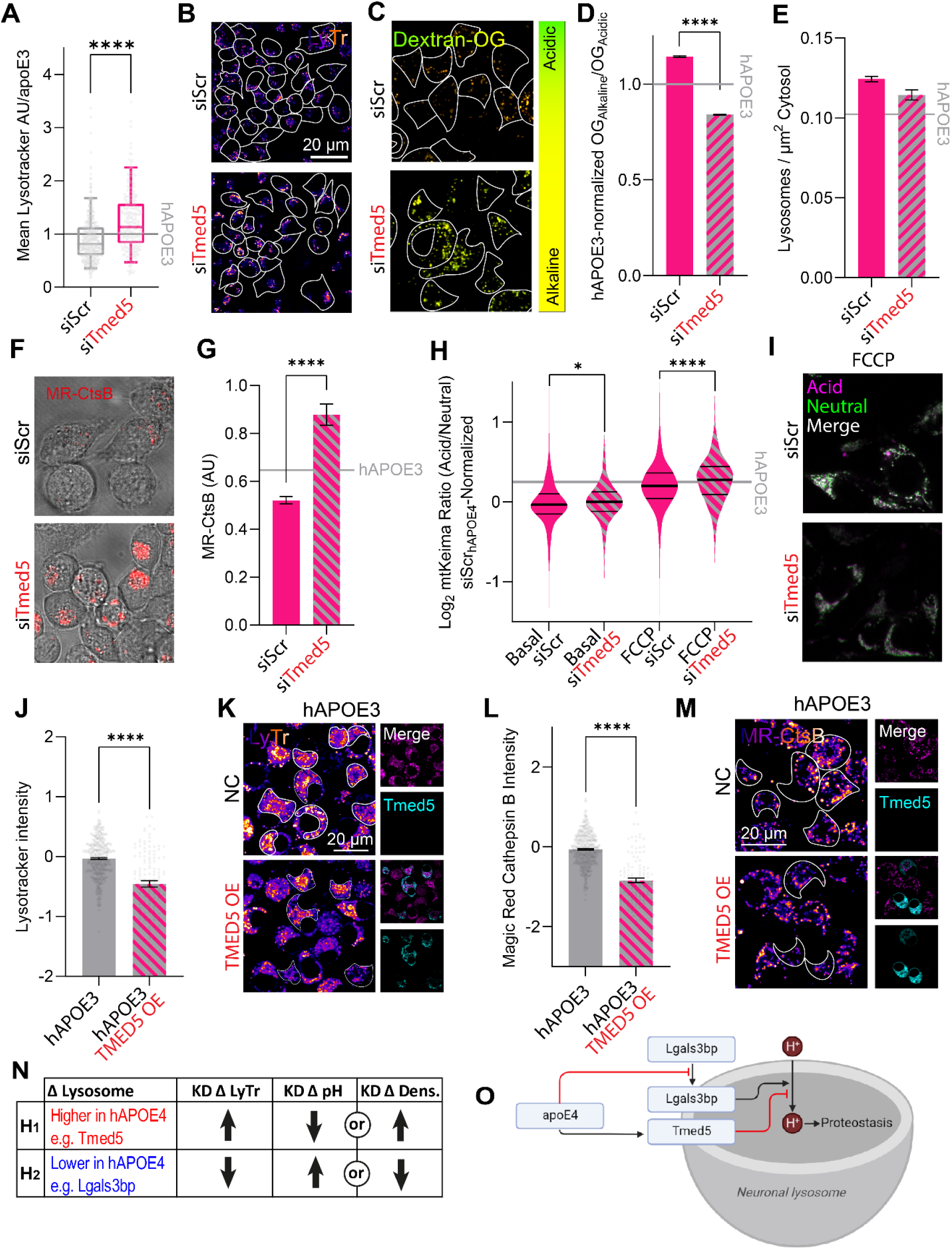
(next page): Lysosomal Tmed5 overflow causes lysosomal alkalinization in hAPOE4 cells. (**A**) Quantification of LysoTracker staining of acidic compartments in hAPOE4 Neuro-2a cells following knockdown of Tmed5. Boxes indicate the 5-95 percentile. Data points indicate individual cells from three independent experiments. (**B**) Representative LysoTracker images for quantification shown in (A). (**C**) Oregon-green relative pH measurements upon Tmed5 knockdown in hAPOE4 Neuro-2a cells, compared to scrambled controls. (**D-E**) Quantification of relative lysosomal pH (D) and density (E) upon Tmed5 knockdown, as shown in panel (C). The value of these measurements in hAPOE3 Neuro-2a cells upon treatment with scrambled controls is shown for reference. Data is plotted as pH per lysosome or cellular lysosomal density across three independent experiments. (**F**) Magic Red Cathepsin B images of hAPOE4 Neuro-2a cells following Tmed5 knockdown. (**G**) Quantification of Magic Red Cathepsin B intensity per cell across three independent experiments, as shown in panel (F). The value of this measurement in hAPOE3 Neuro-2a cells upon treatment with scrambled controls is shown for reference. (**H**) Quantification of mitophagy by mtKeima measurements following Tmed5 knockdown in hAPOE4 Neuro-2a cells under basal and FCCP-induced conditions, plotted as a density plot of mitochondria across three independent experiments. The value of this measurement in hAPOE3 Neuro-2a cells upon treatment with scrambled controls is shown for reference. (**I**) Representative images for panel (H). (**J**) Quantification of LysoTracker staining of acidic compartments in hAPOE3 Neuro-2a cells following human TMED5-GFP overexpression, Log2-normalized relative to untransfected negative controls (NC). Data points indicate individual cells from three independent experiments. (**K**) Representative LysoTracker images for quantification shown in (J). (**L**) Quantification of Magic Red Cathepsin B proteolysis in hAPOE3 Neuro-2a cells following human TMED5-GFP overexpression, Log2-normalized relative to untransfected hAPOE3 cells. Data points indicate individual cells from three independent experiments. (**M**) Representative Magic Red Cathepsin B images for quantification shown in (L). (**N**) Tabular representation of hypotheses. (**O**) Graphical representation of experimental results. All experiments were performed as three technical and biological replicates. Fluorophore intensities and spot counting was performed in Fiji. Groups of data were analyzed in GraphPad Prism by 2-way ANOVA followed *post-hoc* by Dunnett’s multiple comparisons test. Pairs of data were analyzed by t-tests. *P<0.05, ****P<0.0001.

Taken together, we have identified two proteomic driving forces for apoE4-associated lysosomal dysfunction: On one hand, Tmed5 accumulates in apoE4 lysosomes, with its knockdown restoring lysosomal acidity. In contrast, Lgals3bp is depleted from apoE4 lysosomes, and its silencing in apoE3-expressing cells reduces the cellular lysosomal capacity (Fig. 5N). Changes to these lysosomal proteins thereby connect apoE4 expression with the observed lysosomal impairments (Fig. 5O).

### ApoE4 consistently modifies neuronal lysosomes across model systems

Having observed that apoE4 expression impacts lysosomal function through Lgals3bp depletion and Tmed5 accumulation, we evaluated their pathophysiological relevance in human iPSC-derived neurons, neurons from *post-mortem* AD tissue, and apoE4 AD brains as a whole. In order to first obtain cell-type resolution of lysosomal protein accumulation, we used isogenic apoE3 and apoE4 iPSCs to generate mixed neuronal cultures including GABAergic interneurons, which are intimately linked to apoE4 neuropathology [24] (Fig. S7A). We profiled changes in lysosomal LGALS3BP and TMED5 levels using a repurposed proximity ligation assay (PLA), illustrated by Fig. 6A. PLAs are widely used to profile protein-protein interactions, yielding fluorescent signals when the donor and acceptor proteins are within 40 nm of each other [88]. However, the lysosomal lumen is a compact and protein-dense compartment, with a radius ranging from 100-300 nm [89, 90]. We reasoned that coincidental interactions between the reference lysosomal protein LAMP1 and our proteins of interest would occur more frequently in the lysosomal lumen and be detectable by proximity ligation. In agreement with this hypothesis, we found that the LAMP1-LAMP1 positive control yielded a particularly good signal-to-noise ratio over the single-antibody controls, whereas the LAMP1-LGALS3BP and LAMP1-TMED5 yielded comparatively less efficient, yet still significant, lysosomal PLA signals (Fig. S7B-C). Having established that we could detect lysosomal LGALS3BP and TMED5, we compared lysosomal PLA signals between apoE3 and apoE4 human iPSC-derived neurons. In concordance with our observations in Neuro-2a cells, we found that apoE4 iPSC-derived neurons harbored less lysosomal LGALS3BP and more lysosomal TMED5 compared to their apoE3 counterparts (Fig. 6B-C). We subsequently utilized the same assay to assess whether these lysosomal protein changes also occur in *post-mortem* brain slices from AD patients. For LGALS3BP, the PLA signal was generally low, perhaps indicative that its lysosomal depletion could be a wider phenomenon occurring in late-stage AD independently of the apoE4 genotype. While we observed a general trend of lower lysosomal LGALS3BP levels in apoE4 neurons, this was not statistically significant (P=0.24; Fig. 6D-E). On the other hand, we found that neurons from apoE4 AD brain samples accumulate lysosomal TMED5, indicating that lysosomal TMED5 accumulation is a highly conserved event in apoE4 neuropathology (Fig. 6F-G).

**Figure 6.**
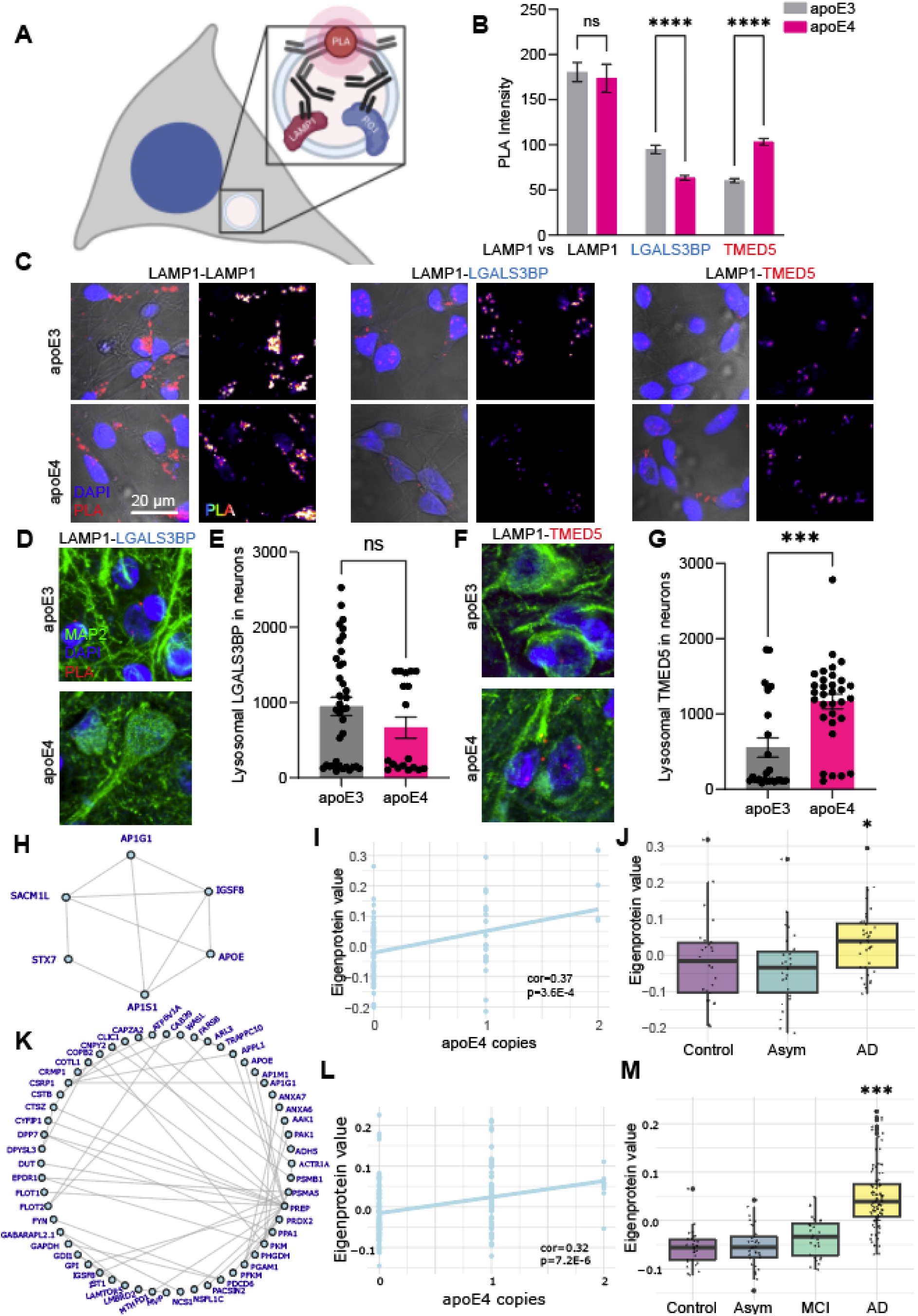
(previous page): ApoE4-associated lysosomal protein changes occur in human AD brains and neurons. (**A**) Schematic of lysosomal proximity ligation assay, whereby proximity ligation occurs between luminal lysosomal LAMP1 and the profiled protein of interest (P.O.I). (**B**) Quantification of cellular proximity ligation assay (PLA) signal in iPSC-derived neurons across three independent differentiations and experiments. LAMP1-LAMP1 was used as a positive control, while lysosomal LGALS3BP and TMED5 localization profiled by PLA with LAMP1. (**C**) Representative images for panel B. (**D**) Representative images of neuronal LGALS3BP-LAMP1 PLA in the AD brain. (**E**) Quantification of lysosomal LGALS3BP in the AD brain across two different cases per genotype, analyzed on a cellular level within a lysosomal mask based on PLA signal and LAMP1 immunoreactivity. (**F**) Representative images of neuronal TMED5-LAMP1 PLA in the AD brain. (**G**) Quantification of lysosomal TMED5 in the AD brain across two different cases per genotype, analyzed on a cellular level within a lysosomal mask based on PLA signal and LAMP1 immunoreactivity. (**H-J**) Plots showing apoE4 lysosome-associated proteins for individual patients from the BLSA cohort, showing (**H**) co-expression (nodes show dysregulated lysosomal proteins, edges denote highest protein co-expression), and their association with (I) APOE4 dosage and (**J**) AD diagnosis. (**K-M**) Plots showing apoE4-dysregulated lysosomal proteins for individual patients from the Banner cohort, showing (**K**) co-expression (nodes show dysregulated lysosomal proteins, edges denote highest protein co-expression), and their association with (**L**) APOE4 dosage and (**M**) AD diagnosis. *P<0.05; ***P<0.0005; ****P<0.0001.

We finally assessed whether the implicated apoE4-dysregulated lysosomal proteins from our LysoIP experiments are conserved across large-scale studies using proteomic datasets from the Accelerating Medicines Partnership – Alzheimer Disease (AMP-AD) consortium [91]. In analogy with our observations that hAPOE4-encoding cells accumulate lysosomal proteins, we first investigated how AD progression affects the lysosomal proteins we identified. Proteomic data from the Baltimore Longitudinal Study of Aging (BLSA) cohort revealed a significant positive correlation between apoE4 lysosome-associated proteins (Fig. 6H) with APOE4 dosage (Fig. 6I, cor=0.37, p=3.6E-4). The same group of proteins were also significantly increased in AD vs controls (Fig. 6J, p=0.03). Moreover, the Banner Sun Health Research Institute (Banner) cohort revealed significant positive correlation between all apoE4-dysregulated lysosomal proteins relative to apoE3 controls (Fig. 6K) with APOE4 dosage (Fig. 6L, cor=0.32, p=7.2E-6). These dysregulated proteins similarly accumulated in AD patients compared to control and asymptomatic, mild-cognitive impairment (MCI) patients (Fig. 6M, AD vs controls p=8.0E-16). Altogether, this data suggest that dysregulated lysosomal proteins may play an important role in the human brain of APOE4 carriers.

These results illustrate that the apoE4-associated and functionally relevant lysosomal protein changes also manifest in human postmitotic neurons susceptible to neurodegeneration. Overall, these findings confirm that apoE4-associated lysosomal protein dysregulation occurs widely and consistently across AD model systems, including cell lines, iPSC-derived human neurons, and *post-mortem* brain samples.

## Discussion

Regardless of its etiology, Alzheimer Disease is riddled with signs of impaired lysosomal function: In healthy neurons, lysosomes clear β-amyloid [92, 93], phospho-Tau aggregates [94], and dysfunctional organelles [95]. In the AD brain however, lysosomal dysfunction drives neuronal accumulation of AD-associated protein aggregates and defective organelles [96]. We find that neuronally encoded apoE4, the primary genetic risk factor for AD, impairs lysosomal function. It is established that astrocytes which express high apoE4 levels suffer from lysosomal alkalinization by unknown mechanisms [97]. While neuronal cells express less apoE4 than astrocytes, we observe a substantial degree of lysosomal alkalinization, which impairs both lysosomal proteolysis and mitophagy.

Focusing on the classical AD hallmark proteins App (forming toxic β-amyloid and β-CTF), we found that hAPOE4 lysosomes to accumulate App and the App-processing γ-secretase Psen2 subunit. This observation suggests that lysosomal defects could affect processing and aggregation of App. Indeed, it has previously shown that endocytic compartments are crucial in the amyloidogenic processing of App [98]. Intriguingly, we found that silencing *App* or its associated *Psen2* did not affect lysosomal function in hAPOE4-expressing cells, suggesting that rather than impairing lysosomal function itself, App accumulation may be downstream of hAPOE4-associated lysosomal impairments.

Our exploration of proteomic alterations to hAPOE4 lysosomes furthermore revealed that hAPOE4 lysosomes carry higher levels of lysosome- and Golgi-associated proteins. We furthermore observed that a select few of these functionally relevant protein changes also manifest in an apoE4 dose-dependent manner in both iPSC-derived human neurons and in AD *post mortem* brain samples, indicating the broader pathophysiological relevance of these observations. In line with previous studies [56, 99], we found hAPOE4 to preferentially localize in lysosomes relative to its hAPOE3 counterpart. This has been postulated as a route for hAPOE4 to escape the secretory pathway, promoting its membrane interaction, cytosolic entry, and subsequent interference with intracellular processes [99–101].

Intriguingly, hAPOE4 expression causes lysosomal accumulation of several Golgi-associated Transmembrane emp24 domain-containing (Tmed) proteins, including Tmed3, Tmed4, Tmed5, and Tmed9. Numerous Tmed proteins have been shown to be dysregulated in mild cognitive impairment (MCI) and AD brains: While TMED5 expression increases in the AD parietal cortex [102], expression of both TMED4 and TMED9 decreases in the AD frontal cortex [103–105]. Tmed proteins regulate processing of secretory proteins and have been primarily studied in the context of coatomer (COP) cargo selection and vesicle formation. However, Tmed proteins also have roles in later secretory compartments [106]. In particular, Tmed5 and Tmed9 have been implicated in trafficking misfolded GPI-anchored proteins from the plasma membrane to the lysosome, alongside Tmed2 and Tmed10 [64]. In line with our own findings, knockdown of TMED2 and TMED10 in human iPSC-derived neurons results in increased lysotracker staining [79], highlighting that the coupling of TMED-family protein expression and lysosomal function is a widely conserved phenomenon.

There is growing evidence that Lgals3bp may have diverse functional roles. Lgals3bp has been studied in the context of cancer biology, and more recently, its neurological involvement has been described in the process of neurodevelopmental corticogenesis. In concordance to our findings in hAPOE4 lysosomes, Lgals3bp expression appears decreased post-in mortem in the Alzheimer Disease entorhinal cortex [107]. While its lysosomal function remains uncharacterized, recent proteomics experiments have demonstrated this protein to be consistently recovered from lysosomes across cell lines [86]. Our results support these previous findings of Lgals3bp as a lysosomal protein and provide insights into how it affects lysosomal function. Specifically, we genetically decreased Lgals3bp expression in hAPOE3 cells, finding that Lgals3bp maintains lysosomal density, and to a lesser extent lysosomal acidity. As Lgals3bp has been localized to extracellular vesicles that form in late endosomal or lysosomal compartments [108] and suppresses endosomal β-amyloid production [109], it is conceivable that Lgals3bp mediates lipid sequestration and fine-tunes β-amyloid generation to prevent β-amyloid-mediated lysosomal damage. We furthermore find Lgals3bp to interact with several subunits of the ER-associated degradation (ERAD) machinery, which routes misfolded proteins from the ER towards lysosomes. We show that hAPOE4 particularly impairs the interaction between Lgals3bp and the ERAD-associated E3 ubiquitin ligase-adaptor Faf2/Ubxd8 through decreasing Faf2 expression [110]. Decreased Faf2 expression has also been shown to facilitate tau seeding [111], indicating that apoE4-sensitive ERAD proteins could be drivers for a host of AD-associated proteostatic and neuropathologic events.

Importantly, while we find that while lysosomal Lgals3bp and Tmed5 are both differentially regulated by apoE4 expression, their impact on lysosomal function is not limited to the context of apoE4. Instead, increased Lgals3bp and decreased Tmed5 may be characteristic of lysosomal health and capable of regulating lysosomal function. This is exemplified by targeted Lgals3bp knockdown resulting in decreased lysosomal density and acidification in apoE3-expressing cells, and CRISPRi-mediated knockdown of several TMED-proteins increasing lysotracker staining in wild-type iPSC-derived neurons [79].

Taken together, we show that lysosomal dysfunction is not limited to early-onset forms of Alzheimer Disease, but also occurs in the more common late-onset disease variant. We show that apoE4 expression impairs lysosomal function in neuronal cells, describe novel mediators of lysosomal pH homeostasis, and highlight their involvement in AD pathophysiology in patients and across experimental models. Lysosomal pH dysregulation occurs in a number of diseases beyond Alzheimer Disease including Parkinson’s Disease, lupus, infectious diseases, cancers, and diabetes, and a better understanding of the processes underlying these phenomena bears promise of broad translational impact [112]. Our findings also implicate apoE4 and tau accumulation as a possible contributor to lysosomal dysfunction in neurons, and provide a rationale for expanding therapeutic trials targeting the lysosomal system to late-onset Alzheimer Disease.

## Materials and Methods

### Plasmids

The vector encoding the lysosomal calcium sensor LAMP2-GCaMP6s (pBoBi-hLAMP2-C-GC6s) was obtained from Addgene (Addgene #154151). The lentiviral packaging vector pΔ-NRF Gag-Pol-Tat-Rev and the pseudotyping vector pMD2.G VSV-G were kindly provided by Prof. Judd Franklin Hultquist of Northwestern University. The lentiviral vector used for lysosomal immunoprecipitation pLJC5-TMEM192-3xHA (Addgene #102930)[63] was kindly provided by Dr. Ali Ghoochani and Prof. Monther Abu-Remaileh of Stanford University. The pCAGGS mtKeima construct [113] was kindly provided by Dr. Huihui Li and Prof. Ken Nakamura of the Gladstone Institutes. The mouse Lgals3bp-GFP construct was cloned by Gibson Assembly. Mouse Lgals3bp was cloned from Neuro-2a cDNA using Q5 polymerase following the manufacturer’s instructions, annealing at 61.2°C with the forward primer caagcttcgaattcagggacATGGCTCTCCTGTGGCTC and the reverse primer cctgtggagccggtggagccCACCATGTCAGTGGAGTTAG. The pEGFP-N3 backbone was subcloned from the LAMP1-mGFP plasmid (Addgene #34831) [114] using the forward primer GGCTCCACCGGCTCCACA and the reverse primer GTCCCTGAATTCGAAGCTTGAGCTC. Gibson Assembly was completed using the NEBuilder HiFi DNA Assembly Master Mix (NEB # E2621L), following the manufacturer’s instructions.

### Neuro-2a Cell Culture

The murine, male neuroblastoma cell line Neuro-2a was previously modified by introducing transgenic, full-length human apoE3 or apoE4 (herein referred to as hAPOE3 and hAPOE4, respectively) [115]. The parental cell line without human apoE expression is referred to as hAPOE^−/−^, referring to its human apoE genotype. Notably, the murine *Apoe* gene is intact in these cells. Neuro-2a cells were cultured in sterile-filtered minimum essential medium + GlutaMAX (MEM, ThermoFisher #41090-036) supplemented with 10% fetal bovine serum (FBS, ThermoFisher #A31605-02), 1X non-essential amino acids (NEAA, ThermoFisher #11140050), and 1 mM sodium pyruvate (ThermoFisher, #11360070). Cells were passaged using Accutase (Fisher Scientific, #NC9839010) and used between passages 5-10.

### Lipofectamine Transfection

Cells were transfected using Lipofectamine 2000 (ThermoFisher #11668019) following the manufacturer’s protocols. Cells were cultured to 70% confluency prior to transfection. For transient transfection, DNA was diluted in OptiMEM to 32 ng/µL and gently mixed. Lipofectamine 2000 was similarly diluted in OptiMEM to 8%, and both mixtures incubated for 5 minutes at room temperature. Following the 5 minutes, DNA-OptiMEM and Lipofectamine-OptiMEM mixtures were combined 1:1, gently mixed, and incubated for 20 minutes at room temperature. The transfection mixture was subsequently added to the cells at a 1:6 mixture-to-media ratio, and the cells transfected overnight. The media was replaced the following day, and the cells were subsequently used for experiments. For siRNA knockdown, we targeted the following genes with the indicated siRNAs (available from ThermoFisher): APOE (AM16708_41598), *App* (s62516), *Lgals3bp* (AM16708_162469), *Mapt* (s70123), *Psen2* (AM16708_68876), scramble control (4390844), and *Tmed5* (AM16703_83238). siRNAs were diluted in OptiMEM to 0.5 µM, and Lipofectamine RNAi Max (ThermoFisher #13778150) diluted in OptiMEM to 5%. siRNA and Lipofectamine mixtures were left for 5 minutes before being combined, and incubated for 20 minutes at room temperature. Following incubation, transfection mixtures were added to the wells at a final concentration of 10% v/v. The cells were first transfected overnight, followed by a complete medium exchange and a second subsequent transfection. The cells were used for experiments after 48 hours knockdown in total.

### RNA extraction, cDNA synthesis and RT-qPCR

We used RT-qPCR to quantify the knockdown efficiencies of our siRNAs. Following 48 hours knockdown, RNA was extracted using the RNeasy Kit (Qiagen #74106), following manufacturers’ instructions, supplementing RLT buffer with β-mercaptoethanol, homogenizing samples by vortexing and without performing gDNA digestion. cDNA was reverse transcribed using the Superscript IV kit (ThermoFisher #18090010), using 1 µg RNA as template, a 0.5:0.5 mixture of Oligo d(T) and random hexamers for priming. The reaction was performed as follows: 10 minutes hexamer annealing at 23°C, 10 minutes amplification at 55°C, and 10 minutes inactivation at 80°C. Reverse-transcribed cDNA was diluted 1:10 with water. RT-qPCR was run using Power SYBR Green PCR Master Mix (Thermo Fisher #4367659) to detect APOE as previously described [116], and TaqMan (ThermoFisher #4444963) to detect *App* (Mm01344172_m1), *Hprt* (housekeeping control; Mm01545399_m1), *Lgals3bp* (Mm00478303_m1), *Mapt* (Mm00521988_m1), *Psen2* (Mm00448405_m1), and *Tmed5* (Mm00547008_m1). RT-qPCR reactions were performed on a CFX Opus 96 real-time PCR system (BioRad) with the following cycling parameters: [95°C, 10:00],40x[95°C, 0:15; 60°C, 1:00] for SYBR reactions, [50°C, 2:00; 95°C, 10:00],40x[95°C, 0:15; 60°C, 1:00] for TaqMan reactions. Expression levels were analyzed by the 2^-ΔΔCt^ approach, using *Hprt* as a housekeeping gene and scramble control samples as the treatment control.

### Immunofluorescence and Confocal Microscopy

Neuro-2a cells were seeded onto poly-L-lysine-coated (Sigma #P4707) 12 mm glass coverslips for confocal microscopy and cultured as previously described. Cells were washed once with PBS (Corning #21-030-CV) before they were fixed with 4% paraformaldehyde (Sigma #F8775). For lysosomal immunostaining, cells were washed three times with PBS, and simultaneously permeabilized and blocked for 1 hour at room temperature in a saponin-based blocking buffer (BB-S; PBS with 1% BSA, 0.05% Saponin, 50 mM NH_4_Cl, and 1% NaN_3_). The cells were stained with mouse anti-HA antibodies (BioLegend #901501) diluted 1:1000, rat anti-mouse Lamp1 antibodies (SantaCruz #sc-19992) diluted 1:100, mouse anti-human LAMP1 antibodies (SantaCruz #sc-20011) diluted 1:800, or rabbit anti-phospho-Tau(S202/T205) (ABclonal #AP0894) diluted 1:1000 in BB-S overnight at 4°C, washed three times with PBS, and stained with either anti-mouse Alexa488-(ThermoFisher #A11001)-, anti-rabbit Alexa488 (ThermoFisher #A21206)-, anti-mouse Alexa555 (#A31570)-, anti-rabbit Alexa555 (ThermoFisher #A21428)-, or anti-rat fluorescein (Vector #FI-4000)-conjugated secondary antibodies diluted 1:500 in BB-S for 2 hours at room temperature. For iPSC-derived neuron immunostaining, cells were similarly fixed in 4% PFA for 15 minutes, and washed once with PBS. Cere washed once with washing buffer (PBS with 0.1% Tween-20), and blocked with blocking solution (PBS with 10% donkey serum and 0.5% Triton-X) for 1 hour. Cells were stained overnight at 4°C with rabbit anti-GABA (ThermoFisher #PA5-32241) diluted 1:1000 and mouse anti-TuJ (Promega #G712A) diluted 1:1000 in PBS with 1% donkey serum. Cells were washed twice with washing buffer, and stained with aforementioned secondary antibodies for 1 hour in PBS with 1% donkey serum. Cells were again washed twice with washing buffer. Nuclei were subsequently stained for by incubation with 200 ng/mL Hoechst 33342 (ThermoFisher #H3570) diluted in PBS for 30 minutes at room temperature. The samples were washed three times with PBS and mounted onto Micro Slides (VWR #48311-703) using Cytoseal™ 60 (VWR #8310-4). Images were captured using a confocal laser scanning microscope (Zeiss LSM880) fitted with a Plan-Apochromat 63x/1.4 Oil DIC M27 objective and PMT detector. Nuclei were imaged by excitation at 405 nm and detection between 410-507 nm, with the pinhole set to 1.36 AU. The AlexaFluor488-stained lysosomal tags were imaged by excitation at 488 nm and detection between 493-630 nm, with the pinhole set to 0.43 AU. Pixel scaling was set to 70 nm to capture lysosomal structures.

### Lysotracker Imaging

The lysosomal expanse of the Neuro-2a cells was determined using the Lysotracker probe. Neuro-2a cells were seeded into live-cell imaging chambers (ibidi #80806) at a density of 2*10^5^ cells/mL. The media was exchanged for fresh media on day 2 of culture. On day 3, the cells were stained with Lysotracker Red DND-99 (ThermoFisher #L7528) diluted 1:10,000 to label acidic endolysosomes and CellMask™ Deep Red Plasma Membrane Stain (ThermoFisher #C10046) diluted 1:1000 to label the plasma membrane. The cells were loaded for 30 minutes at 37°C before being chased in culture media not containing staining reagents. The cells were transferred to a confocal laser scanning microscope (Zeiss LSM880) fitted with a Plan-Apochromat 63x/1.4 Oil DIC M27 objective and PMT detector, with its incubation chamber pre-heated to 37°C and atmosphere filled with 5% CO_2_. Lysotracker staining was imaged by excitation at 561 nm and detection between 566-639 nm, with the pinhole set to 0.81 AU. CellMask staining was imaged by excitation at 633 nm and detection between 639-759 nm, with the pinhole set to 0.93 AU. Pixel scaling was set to 90 nm to capture lysosomal structures. Image analysis was performed in Fiji. The CellMask channel was used to manually draw regions of interest (ROIs) delimiting individual cells, and the roiManager Measure function used to measure the Lysotracker staining intensity of each cell. The results shown were obtained from three distinct biological and technical replicate experiments.

### pH Measurements using Oregon Green

The lysosomal pH was accurately determined using the Oregon Green probe. Neuro-2a cells were seeded into live-cell imaging chambers (ibidi #80806) at a density of 2*10^5^ cells/mL. On day 3, the cells were loaded with 500 µg/mL OregonGreen™ 488; Dextran 70,000 MW (ThermoFisher #D7172) overnight. The following day, the cells were chased in culture medium for 1 hour, before 0.1% DMSO (Sigma #D2650) or 200 nM Bafilomycin A1 (FisherScientific #AAJ61835MCR) was added for another hour. The cells were transferred to a confocal laser scanning microscope (Zeiss LSM880) fitted with a Plan-Apochromat 63x/1.4 Oil DIC M27 objective and PMT detector. pH-insensitive, reference OregonGreen™ 488 staining was imaged by excitation at 458 nm and detection between 495-605 nm. pH-sensitive OregonGreen™ 488 staining was imaged by excitation at 488 nm and detection between 495-605 nm. The pinhole was set to 1.14 AU, and pixel scaling set to 135 nm to capture lysosomal structures. After capturing 7 images of each condition, the lysosomes were pH-clamped using the Intracellular pH Calibration Buffer Kit (ThermoFisher #P35379), with its range expanded to include 10 different pH values ranging from pH 3.60 to pH 7.50, adjusting the buffer pH with HCl. The buffer kit was used following the manufacturer’s instructions, adding buffers starting at pH 7.50 and progressively adding more acidic buffers. One calibration image was captured for each pH buffer. Image analysis was performed in Fiji. The CellMask channel was used to manually draw single regions of interest (ROIs) delimiting cells. A mask was generated of endosomes by using the 456 nm reference channel. A minimum filter was first applied followed by a maximum filter (radius=3). The Li AutoThreshold function was used to create a binary mask of lysosomes, followed by a dilation of the cell mask using EDM Binary Operations (iterations=3). The Watershed function was used to divide lysosome clusters, and the mask used to create a region of interest. The region of interest as split into its individual constituents, and the intensity of each individual ROI (endosome) measured from the raw, unprocessed image. The ratio of pH-sensitive to pH-insensitive fluorescence was subsequently calculated for each individual endosome. The same data analysis was performed for the pH-clamped lysosomes, allowing extrapolation from Oregon Green ratios to their respective absolute pH values. The results shown were obtained from three distinct biological and technical replicate experiments.

### Calcium Imaging using LAMP2-GCaMP6s

Neuro-2a cells were seeded into live-cell imaging chambers (ibidi #80806) and cells cultured to confluence. Upon reaching confluence, the cells were transiently transfected for 48 hours with the LAMP2-GCaMP6s construct [60] using Lipofectamine 2000 (ThermoFisher #11668019) following the manufacturer’s instructions. Following transgene expression, the medium was replaced with Ca^2+^-free standard bath solution (142 mM NaCl, 6 mM KCl, 2 mM MgCl_2_, 5.5 mM Glucose, 1 mM EGTA, 10 mM HEPES, pH 7.4). Cells were transferred to a Zeiss LSM880 confocal microscope fitted with a Plan-Apochromat 63x/1.4 Oil DIC M27 objective and an Airyscan detector, preequilibrated to 37°C and 5% CO_2_. Calcium-bound GCaMP6s was imaged by excitation at 488 nm and detection between 415-735 nm. To facilitate high-speed imaging while maintaining resolution of lysosomal structures, pixel scaling was set to 130 nm, the pinhole set to 2.31 AU, and images captured every second for 250 seconds. Cells were treated with 200 µM GPN (Cayman #14634) at 50 seconds, and with 4 µM Ionomycin (Cayman #10004974) at 150 seconds. At least three movies of were captured for each technical replicate. Image analysis was performed in Fiji. A region of interest was generated of transfected cells excluding the nuclei, delimiting the area to be analyzed. The fluorescence intensity of each region of interest (cell) was measured over time using the Fiji MultiMeasure function. The results shown were obtained from three distinct biological and technical replicate experiments.

### Mitophagy Measurements using mtKeima

Neuro-2a cells were seeded into live-cell imaging chambers (ibidi #80806) at a density of 2*10^5^ cells/mL, and cells cultured to 70% confluence. Upon reaching 70% confluence, the cells were transfected with the mtKeima construct[113, 117] using Lipofectamine 2000 (ThermoFisher #11668019) following the manufacturer’s instructions. The cells were transfected overnight, and media changed the next day. Fresh media was added alongside compounds for treatment, including 0.1% DMSO, 10 µM FCCP (Sigma #SML2959), and 200 nM Bafilomycin A1. The cells were incubated for 30 minutes prior to imaging. The cells were transferred to a confocal laser scanning microscope (Zeiss LSM880) fitted with a Plan-Apochromat 63x/1.4 Oil DIC M27 objective and PMT detector, with its incubation chamber pre-heated to 37°C and atmosphere filled with 5% CO_2_. Neutral mtKeima was imaged by excitation at 488 nm and detection between 600-708 nm, while simultaneously capturing transmitted light. Acidic mtKeima was imaged by excitation at 514 nm and detection between 544-680 nm. The pinhole was set to 0.92 AU, and pixel scaling set to 90 nm to capture mitochondrial structures. Five images were captured for each condition. Image analysis was performed in Fiji. A region of interest was generated of transfected cells based on the transmitted light, delimiting the area to be analyzed. A mask was generated of mitochondria by using both mtKeima channels. A minimum filter was first applied followed by a maximum filter (radius=3). The Li AutoThreshold function was used to create a binary mask of lysosomes, followed by a dilation of the cell mask using EDM Binary Operations (iterations=3), watershed, and erosion using EDM Binary Operations (iterations=3). The resulting mask was used to create a region of interest. The region of interest as split into its individual constituents, and the intensity of each individual ROI (mitochondrion) measured from the raw, unprocessed image. The ratio of acidic to neutral mtKeima was calculated for each mitochondrion. The results shown were obtained from three distinct biological and technical replicate experiments.

### Lentiviral Production

Lentiviruses were produced as previously described [118]. HEK293T were cultured in DMEM (Corning #10-017-CV) supplemented with 10% fetal bovine serum (FBS, ThermoFisher #A31605-02), 1 mM sodium pyruvate (ThermoFisher, #11360070), and 1X Penicillin-Streptomycin (Corning, #30-002-CI), and used between passages 4-8. On day 1, 6*10^6^ cells were seeded in 50 mL culture medium in 15 cm dishes and allowed to adhere overnight. On day 2, transfection mixtures were prepared by diluting 5 µg lentiviral delivery vector, 3.33 µg Gag-Pol-Tat-Rev packaging construct, and 1.66 µg VSV-G envelope construct in 250 µL serum-free DMEM (Corning #10-017-CV). 30 µL PolyJet In Vitro DNA Transfection Reagent (SignaGen #SL100688) was similarly diluted in 250 µL serum-free DMEM, and incubated at room temperature for 5 minutes. Following the 5 minutes, DNA-DMEM and PolyJet-DMEM mixtures were combined 1:1, gently mixed, and incubated for 25 minutes at room temperature. The transfection mixture was subsequently added dropwise to the cells, plates swirled, and incubated for 72 hours. The supernatant was harvested on day 5. The supernatant was collected, and cellular debris removed by centrifugation at 400g for 5 minutes. The supernatant was subsequently filtered using a 0.45 µm PVDF filter unit, and virions precipitated by adding 8.5% PEG-6000 and 0.3M NaCl followed by 2 hours incubation at 4°C. Virions were pelleted by centrifugation at 2700 g for 20 minutes at 4°C. The supernatant was decanted, virions resuspended in 250 µL PBS, and aliquoted for long-term storage at −80°C.

### Lentiviral Transduction

Neuro-2a cells were transduced by lentiviral spinoculation. Neuro-2a cells were dissociated using Accutase (Fisher Scientific, #NC9839010), and 3*10^4^ cells resuspended in 2 mL media. Polybrene (Millipore #TR-1003-G) was added to the cells to a final concentration of 8 µg/mL alongside 100 µL concentrated lentivirus, and the cells centrifuged at 1000 g for 1 hour at room temperature. The supernatant was aspirated, cells resuspended in 2 mL Neuro-2a media (described in Neuro-2a Cell Culture section) and seeded in a single well of a 6-well tissue culture plate. The cells were cultured for 48 hours post-transduction before selecting for transduced cells using 2 µg/mL Puromycin (ThermoFisher #A1113802) for 3 days. Following antibiotic selection, the polyclonal cells were passaged and expanded once before being frozen for further experiments.

### Lysosomal Immunoprecipitation

Lysosomal immunoprecipitations were performed in biological and technical triplicates as performed as previously described [63] with minor alterations. Neuro-2a cells with each combination of apoE genotype (hAPOE^−/−^, hAPOE3, or hAPOE4) and lysosomal tag integration (background or LysoIP-tagged) were seeded, with 2*10^6^ cells being seeded across two 15 cm dishes per replicate. To distinguish intrinsically lysosomal proteins from lysosomal cargo destined for degradation, two 15 cm dishes of LysoIP-tagged hAPOE^−/−^, hAPOE3, and hAPOE4 Neuro-2a cells were treated with 200 nM Bafilomycin A1 (ThermoFisher #AAJ61835MCR) for 3 hours, preventing lysosomal degradation and enriching for lysosomal cargo. To harvest lysosomes, each sample was individually processed from beginning to lysis. All following steps were performed on ice or at 4°C ambient temperature, unless otherwise stated. Magnetic anti-HA beads (ThermoFisher #88837) were washed twice with KPBS (136 mM KCl, 10 mM KH_2_PO_4_, pH adjusted to 7.25). Culture media was quickly decanted and rinsed twice with ice-cold KPBS. Cells were collected by scraping and transferred to microfuge tubes. Cells were pelleted by centrifugation (1000g, 2 min), and the supernatant aspirated. The cells were next resuspended in 950 µL KPBS, and 25 µL cell suspension aliquoted into Protein LoBind tubes for whole-cell analysis. The remaining 925 µL cell suspension was homogenized upon passing the entire volume through a 20G needle 10 times, and efficient release of cellular organelles assessed under a phase contrast microscope. The homogenate was transferred to a new microcentrifuge tube and centrifuged (1000g, 2 min) to pellet the plasma membrane. The supernatant was transferred to a new microcentrifuge tube containing 150 µL magnetic anti-HA beads and mixed once by pipetting. The organelles and beads were rocked at 1200 rpm for 3 minutes and transferred to a magnet for washing. The beads were incubated on the magnet for 25 seconds before the supernatant was aspirated and replaced with fresh KPBS. The beads were rocked at 1200 rpm for 10 seconds and returned to the magnet. These washes were repeated three times, transferring the cells to Protein LoBind tubes for the final wash. Upon aspirating the KPBS following the final wash, 300 µL modified SP3 lysis buffer [119] (50 mM Tris-HCl, 50 mM NaCl, 1% SDS, 1% NP-40, 1% Tween 20, 1% Glycerol, 1% Sodium Deoxycholate, 5 mM EDTA, 5 mM Dithiothreitol, 5 KU Benzonase, 1 Complete protease inhibitor; modified by excluding Triton X-100) was added to the input and bead samples. Upon finishing processing all samples, the samples were lysed by rocking at 65°C for 30 min at 1200 rpm. LysoIP samples were placed on the magnet for 25 seconds, and the supernatant transferred to fresh Protein LoBind tubes to remove magnetic beads. The samples were alkylated by adding chloracetamide (final concentration 10 mM) and shielded from light for 30 minutes. The alkylated samples were vortexed, solubilized on ice for 20 minutes, and centrifuged at 16000g for 10 minutes. The soluble supernatant was transferred to new tubes, and −20°C acetone added in four-fold volume to precipitate protein. The samples were briefly vortexed and incubated at −80°C for 1 hour. Proteins were pelleted by centrifugation (2000 g, 15 min), and the pellet washed twice with −20°C acetone. The samples were stored at −80°C until mass spectrometric analysis.

### Mass Spectrometry

Dry, digested peptides, were resuspended in 0.1% FA and separated using Thermo EASY-nLC 1200 nano liquid chromatography setup. Separation was performed using a 15 cm long Bruker PepSep column with a 150 µm inner diameter packed with 1.5um Reprosil Saphir C18 particles. Mobile phase A was composed of a 0.1% formic acid solution, while mobile phase B was composed of 0.1% formic acid with 80% acetonitrile. Samples were loaded onto the column at the maximum flow rate possible with an upper pressure limit of 300 bar, with the gradient performed using a stable flow of 600 nl/min. For LysoIP samples, mobile phase began at 2%, before increasing to 30% over 70 mins. Mobile phase B then increased to 35% over 8 minutes before increasing to 90% B over 2 minutes and finishing with a wash at 90% B for 10 minutes. The gradient required a total time of 90 minutes. For APMS samples, mobile phase began at 4% B before increasing to 35% over 44 minutes. Mobile phase B then increased to 45% over 5 minutes before increasing to 88% B over 1 minute and finishing with a wash at 88% for 10 mins. The gradient for APMS samples required a total time of 60 minutes.

Eluting peptides were ionized using electrospray ionization from a PepSep stainless steel emitter and analyzed using a Thermo Orbitrap Fusion Lumos mass spectrometer. For LysoIP samples acquisition was performed in a data-independent (DIA) manner with a single survey scan from 350-1200 m/z at 120,000 resolution, followed by 40 variable window (Table S3) MS2 scans at 30,000 resolution with stepped HCD using 32 +/- 5 NCE. Survey scans utilized an AGC target of 4e5 with maximum injection time set to “Auto”, while MS2 scan AGC target was set to 5e4 with a maximum injection time of 54 ms. All scans were taken in the Orbitrap. For APMS samples, acquisition was performed in a data-dependent (DDA) manner. Survey scans were taken from 350-1350 m/z at a resolution of 240,000, with a normalized AGC target of 250%, and a maximum injection time 50 ms. Dependent MS2 scans were collected in the ion trap from 200-1200 m/z using the “rapid” scan speed, with a normalized AGC target of 300%, a “dynamic” maximum injection time, and an isolation window of 0.7 m/z. Ions were activated using HCD with 32 NCE. Only ions with charge state 2-5 were selected for fragmentation and ions were excluded with a 10 ppm tolerance for 20 seconds after being isolated for MS2 scans twice. MS system suitability was monitored with QCloud2 [120].

For APMS samples, raw files were searched in MaxQuant using default parameters against a full reviewed mouse proteome including isoforms (downloaded from Uniprot on June 6, 2022) to which human APOE and TMEM192 proteins were added. The evidence.txt output table from that search was then used to generate the three tables necessary for SAINTexpress scoring (baits, preys, and interactions) using the artmsEvidenceToSaintExpress function from the artMS package (release 3.17) in R (version 4.1.1). Two sets of bait, prey, and interaction tables were generated, one using spectral counts and the other using intensity, with GFP samples set as the control group. Interactions were scored using SAINTexpress (version 3.6.3). Protein interactions were identified as significant if they resulted in a BFDR < 0.05 in either spectral count- or intensity-based SAINT analysis, and the prey protein was observed in all three replicates of that hAPOE-bait group (e.g., all Lgals3bp-GFP/hAPOE3, or all Lgals3bp-GFP/hAPOE4 samples). For these significant interactors we calculated protein fold change and significance when comparing the two alleles (Lgals3bp-GFP/hAPOE3 vs. Lgals3bp-GFP/hAPOE4) using the LFQ intensity from the proteinGroups.txt output table from MaxQuant. The identified Lgals3bp interaction partners and their quantification between apoE alleles are provided in Table S2.

For LysoIP samples, the resulting raw files were searched using directDIA in Spectronaut against a full reviewed mouse proteome including isoforms (downloaded from Uniprot on June 6, 2022) to which human APOE and TMEM192 proteins were added. Default search parameters were used without cross-run normalization or imputation. The resulting MSstats-formatted report was then used to summarize abundance at the protein level using Tukey’s median polish after median normalization of peptide features [121]. Proteins were annotated as being lysosomal if they fulfilled one of two conditions: Either they were detected in three tagged replicates but undetected in untagged controls, or they were more than 2-fold enriched in lysosomes over the background, and reaching statistical significance. If proteins were only passing these criteria in Bafilomycin A1-treated samples, they were annotated as being lysosomal cargo. A subset of correlated peptide ions specific for human TMEM192 were selected for quantification of TMEM192 protein, to allow for the most accurate normalization of protein abundance to lysosomes (Table S1G). The identified lysosomal proteins, their quantification between alleles and the precursors used for normalization are provided in Table S2.

Proteins found to be significantly up- or downregulated in hAPOE4 lysosomes relative to hAPOE3 or hAPOE^−/−^controls were tested for enrichment of Gene Ontology (GO Biological Process, Molecular Function and Cellular Component) terms. The over-representation analysis (ORA) was performed using the enricher function from R package clusterProfiler (version 4.2.2) [122]. The gene ontology terms and annotations were obtained from the R annotation package org.Mm.eg.db (version 3.8.2). Non-redundant GO terms were selected by first constructing a term tree based on distances (1-Jaccard Similarity Coefficients of shared genes in GO database) between the significant terms using the R function hclust. The term tree was then cut at a specific level (R function cutree, h = 0.99) to identify clusters of redundant gene sets. For results with multiple significant terms belonging to the same cluster, we selected the most significant term (i.e., minimum adjusted p-value).

Principal component analysis and significance testing were performed using base R (svd and t.test functions respectively). Select protein heatmaps were generated using pheatmap package. Lysosomal protein and gene ontology heatmaps were generated using the complexHeatmap package [123, 124]. Lysosomal overrepresentation analysis was performed by plotting ratios of ratios. The first ratio is the lysosomal distribution coefficient, calculated as the lysosomal protein abundance relative to the whole-cell protein abundance. Proteins that preferentially localize to lysosomes have a high lysosomal distribution coefficient. Conversely, proteins that prefer cytosolic residency have low lysosomal distribution coefficients. To elucidate whether proteins are overrepresented or underrepresented on hAPOE4 lysosomes, we divided the lysosomal distribution coefficients of each protein on hAPOE4 lysosomes by the respective lysosomal distribution coefficient in control cells.

### Lgals3bp-GFP Affinity Purification

For each replicate, Neuro-2a cells stably expressing either hAPOE3 or hAPOE4 were seeded onto two 15 cm dishes at a density of 400,000 cells, and allowed to grow for 4 days. Each 15 cm dish was transfected with 60 µg DNA using polyJET (SignaGen #SL100688) to deliver expression vectors for GFP or Lgals3bp-GFP, following the manufacturer’s instructions. For each vector, three independent biological replicates were prepared. Media was exchanged the following day, and the cells allowed to recover for another three days before harvesting. The subsequent affinity purification was performed as previously described [31]. For each sample, two 15 cm dishes were each rinsed with PBS and scraped in 500 μL NP40 lysis buffer (50 mM Tris-HCl pH 8.0, 150 mM NaCl, 1 mM EDTA pH 8.0, 0.5% Nonidet P40 Substitute, complete protease inhibitor tablet, phosSTOP phosphatase inhibitor tablet) and combined before freezing on dry ice for 20 minutes. Samples were next thawed in a 37°C water bath for lysis until completely thawed, and frozen at - 80°C until GFP immunoprecipitation. Frozen samples were again partially thawed at 37°C, and incubated at 4°C on a tube rotator for 30 minutes. The debris was pelleted by centrifugation (13,000 g, 4°C, 15 minutes) and the supernatant transferred to a 96-well deep well plate and kept on ice until GFP immunoprecipitation.

In addition to lysates, beads and buffers (indicated below) were dispensed into KingFisher 96-well deep-well plates or microplates as appropriate and placed on ice until loaded onto the KingFisher Flex (KFF) Purification System (Thermo Fisher Scientific) for automated processing as follows: GFP-Trap beads (25 μL slurry dispensed in plate 1 only; Cat. #GTMA-10, Chromotek) were equilibrated twice (plates 1,2) with up to 1.0 mL IP Buffer (50 mM Tris–HCl, pH 7.4 at 4 °C, 150 mM NaCl, 1 mM EDTA) supplemented with 0.05% NP40 and incubated with (plate 3) 1.0 mL cell lysate for 2 h. Protein-bound beads were washed three times (plates 4-6) with 1.0 mL IP Buffer supplemented with 0.05% NP40 and then once (plate 7) with 1.0 mL IP buffer before elution. Proteins were eluted from beads (plate 8) in 50 μL 0.05% RapiGest in IP Buffer and combined with residual proteins recovered by rinsing beads (plate 9) in 50 μL IP Buffer for sample processing (below). The KFF is operated in a cold room to maintain a 4°C temperature during immunoprecipitation; however, elution and the final bead rinsing steps were performed using a heat block pre-heated to 23°C. Automated protocol steps were performed using the slow mix speed and the following mix times: 30 seconds for equilibration/wash steps, 2 hours for binding, 35 minutes for elution and 2 minutes for the final bead rinse. Five 30 second bead collection times were used at the end of each step before transferring beads to the next plate.

Proteins from elution and bead rinse steps were combined in a 96-well PCR plate for sample processing as follows: denaturation and reduction at 37°C for 30 m with 2 M urea and 1 mM DTT in 50 mM Tris–HCl pH 8.0, alkylation at room temperature in the dark for 45 m with 3 mM iodoacetamide, and quenching for 10 minutes with 3 mM DTT. Trypsin (0.5 μg/μL; Promega) was added twice (1.0 μL and 0.5 μl) and incubated at 37°C for 4 hours and 2 hours, respectively. Incubations were performed at 37°C in a thermal cycler or at room temperature in a MixMate incubator with shaking at 1,000 rpm. Peptides were acidified with TFA (0.5% final, pH < 2.0) and desalted at room temperature using a BioPureSPE Mini 96-well plate (20 mg PROTO 300 C18; The Nest Group). Briefly, desalting columns were sequentially equilibrated with 0.2 mL 100% methanol; 0.3 mL 80% ACN, 0.1% TFA and 0.3 mL 2% ACN, 0.1% TFA before passing acidified samples through columns twice and subsequently washed with 2% ACN, 0.1% TFA (0.1 mL and 0.4 mL) and 0.1% FA (0.4 mL, twice). Peptides were eluted twice with 50% ACN, 0.1% FA (60 µL each step) and dried under vacuum centrifugation (CentriVap Concentrator, Labconco). The desalting plate was centrifuged at 2,000 rpm for 2 minutes for initial equilibration steps and 3 minutes for all remaining steps. All buffers were prepared with HPLC or LC-MS grade reagents. Within a respective apoE genotype cell line, proteins found to be consistently detected in all Lgals3bp-GFP replicates and significantly enriched over GFP-transfected controls were considered Lgals3bp interaction partners.

### Small molecule-mediated differentiation of human iPSC-derived neurons

The human female apoE4 iPSCs and the isogenic apoE3 iPSCs were previously described [24]. Human iPSCs were maintained on Matrigel (Corning #354277) in mTeSR Plus Media (StemCell Technologies #100-0274) and passaged into media supplemented with 10 µM Y-27632 ROCK inhibitor (Tocris #1254) using Accutase (Millipore #scr005). Mixed neurons were generated as reported previously[24], with some modifications. Briefly, iPSCs from 6 100 mm dishes were suspended as single cells in 50 mL KSR medium (80% DMEM #10-566-024, 20% KOSR #10828028, 1X NEAA, 1X GlutaMAX, 0.5X Pen/Strep, 10 µM β-mercaptoethanol) supplemented with 10 µM SB-431542 (Stemgent #04-0010-10), 250 nM LDN-193189 (Stemgent #04-0074), 5 µM IWP2 (Millipore #506072), 100 nM SAG (Millipore #566661), and 10 µM ROCK inhibitors and placed into two T75 flasks to form neurospheres. Neurospheres were fed every second day until day in vitro 7 (DIV7), at which point the KSR small molecule supplements were replaced with 100 ng/mL FGF8 (Peprotech #100-25), 5 µM IWP2, 100 nM SAG, and 250 nM LDN-193189. At DIV14, neuroepithelium was plated down onto two poly-L-lysine (Sigma #4707)/Laminin (ThermoFisher #23017-015)-coated 100 mm cell culture dishes in N2 media (100% DMEM/F12 #11330-032, 0.5X N2 Supplement #17502-048, 1X NEAA, 1X GlutaMAX, 0.5X Pen/Strep) supplemented with 100 ng/mL FGF8 and 100 nM SAG. At DIV21, neuroepithelium was passaged with Accutase, strained through 40 µm filters, counted and centrifuged (200g, 2 min), before they were resuspended in N2/B27 medium (50% DMEM/F12, 50% Neurobasal #21103-049, 0.5X N2 Supplement, 0.5X B27 Supplement (ThermoFisher #17504-044), 1X NEAA, 1X GlutaMAX, 0.5X Pen/Strep) supplemented with 10 ng/mL BDNF (Peprotech #450-02) and 10 ng/mL GDNF (Peprotech #450-10) and seeded at a density of 200,000 cells/mL onto PLL/Laminin-coated glass cover-slips. iPSC-derived neurons were fed twice per week by half feeds, and used at DIV49.

### Generation of CRISPRi-repressed iPSC-derived NGN2 neurons

The WTC11 iPSCs with AAVS1-integrated doxycycline-inducible mouse NGN2 and CLYBL-integrated dCas9-BFP-KRAB (CRISPRi) were previously described (CRISPRi-i^3^N iPSCs)[78]. sgRNAs targeting TMED10 ([TGGAGACTCGTTCACCACCGA]), LGALS3BP ([GGCCTGACCACGCTCCATAC]) were cloned into the pMK1334 screening vector and verified by Sanger sequencing. The plasmids were then used to produce lentiviruses, and these used to transduce CRISPRi-i3N iPSCs at 70% MOI. Transduced cells were subsequently selected for my puromycin treatment (1 µg/mL), and following selection cultured in StemFlex Medium (Catalog number: A3349401) on GFR, LDEV-free Matrigel-coated dishes (Corning #356231), diluted 1:100 in Knockout DMEM (Thermo Fisher #10829-018). StemFlex was replaced every day, or every other day once 50% confluent, and passaged upon reaching 80-90% confluence using StemPro Accutase (Thermo Fisher #A11105-01). Single-cell iPSCs were always seeded in StemFlex supplemented with 10 nM Y-27632 ROCK inhibitors (Tocris #125410). To generate glutamatergic neurons, iPSCs were passaged and resuspended at 4 x 106 cells per Matrigel-coated T-75 flask in N2 pre-differentiation medium (100% Knockout DMEM/F12 #12660-012, 1X NEAA, 1X N2 Supplement #17502-048, 10 ng/mL NT-3 #450-03, 10 ng/mL BDNF# 450-02, 1 µg/mL Laminin #23017-015, and 2 µg/mL Doxycycline hydrochloride #D3447-500MG). During plating and for 24 hours only, 10 nM Y-27632 ROCK inhibitor was added to the pre-differentiation medium. After three days at DIV0, cells were passaged with accutase and resuspended at [30,000 cells per well on Corning® BioCoat® Poly-D-Lysine 96-well Black, flat-bottomed plates (354640 in classic neuronal medium (50% DMEM/F12 #11320-033, 50% Neurobasal-A #10888-022, 1X NEAA, 0.5X GlutaMAX, 0.5X N2 Supplement, 0.5X B27 Supplement #17504-044, 10 ng/mL NT-3, 10 ng/mL BDNF, 1 µg/mL Laminin, and 2 µg/mL Doxycycline hydrochloride. On day 7, half of the medium was replaced with fresh classic neuronal medium without doxycycline. On day 14, half of the medium was removed, and twice this amount of fresh neuronal medium was added back to the cells. The cells were imaged by confocal microscopy on day 21.

### Brain Tissue Samples

Human brain tissue samples were acquired from the Neurodegenerative Disease Brain Bank at the University of California San Francisco. Neuropathological diagnoses were made following consensus diagnostic criteria [125]. Cases were selected based on clinical and neuropathological diagnoses, and stratified based on apoE genotype. All *post-mortem* samples were obtained from patients showing AD pathology at Braak stage 6/Thal stage 5. ApoE3 patients showed an age of onset of 55 and 59 years, and age of death of 75 and 76 years, respectively, while apoE4 patients showed an age of onset of 60 and 53 years, and age of death of 66 and 74, respectively. Frozen human brain tissue samples were dissected from the angular gyrus of apoE3 and apoE4 cases.

### Proximity Ligation Assay

The proximity ligation assay (PLA) was performed following the manufacturer’s instructions (Millipore #DUO92101), with minor adaptations to detect lysosomal luminal antigens. For neuronal cultures, DIV49 iPSC-derived neurons were washed twice with calcium-containing PBS, fixed with 4% paraformaldehyde for 30 minutes, and washed once with calcium-containing PBS prior to the assay. For histological samples, paraffin-embedded angular gyrus samples from homozygous apoE3 or apoE4 AD patients were sliced 8 µM thick and dewaxed by baking (65°C, 30 min), xylene (10 min) and ethanol deparaffination (2×5 min 100% ethanol, 1 min 100% ethanol, 1 min 95% ethanol), peroxidase blocking by 30 min 3% H_2_O_2_/methanol incubation, and 0.01M citric acid antigen retrieval at 121°C for 5 minutes. All samples were permeabilized and blocked with BB-S for 1 hour at room temperature. Subsequently, the samples were incubated with the following primary antibodies in BB-S at 4°C overnight: mouse anti-LAMP1 (SantaCruz #sc-20011, diluted 1:800), rabbit anti-LAMP1 (CST #9091S, diluted 1:200), rabbit anti-LGALS3BP (ThermoFisher #10281-1-AP, diluted 1:200), and rabbit anti-TMED5 (ThermoFisher #PA5-141055, diluted 1:500). Histological slides were additionally stained with chicken anti-MAP2 (abcam #ab5392, diluted 1:10,000).The samples were washed twice in home-made wash buffer A to detect lysosomal antigens (10 mM Tris, 150 mM NaCl, 0.05% Saponin) for 5 minutes, and incubated in a humid chamber with PLA probe solution containing 1X PLUS and MINUS PLA probes against mouse and rabbit primary antibodies for 1 hour at 37°C. The samples were washed twice for 5 minutes with lysosomal wash buffer A, and incubated in 1X ligation solution for 30 minutes in a humid chamber at 37°C. The samples were washed twice for 5 minutes with lysosomal wash buffer A, and incubated in 1X amplification solution for 100 minutes in a humid chamber at 37°C. The samples were finally washed twice for 10 minutes in 1X wash buffer B (200 mM Tris, 100 mM NaCl), once for 1 minute in 0.01X wash buffer B, and mounted in Duolink In Situ Mounting Medium with DAPI on glass slides for confocal microscopy. Images were captured using a confocal laser scanning microscope (Zeiss LSM880) fitted with a Plan-Apochromat 63x/1.4 Oil DIC M27 objective and PMT detector. Nuclei were imaged by excitation at 405 nm and detection between 410-507 nm, with the pinhole set to 1.36 AU. PLA signals were captured using TexasRed excitation/emission settings, with the pinhole set to 1.40 AU. Pixel scaling was set to 70 nm. PLA signals were analyzed in Fiji, drawing region-of-interests around neuronal soma based on PMT and DAPI images, followed by average signal intensity measurements. The experiment was performed in triplicate, using neurons from three different differentiations.

### Validation of lysosomal proteomic dysfunction in the postmortem human neurodegenerative disease brain

Label-free quantitative proteomic data from postmortem human brain tissues were downloaded for the Baltimore Longitudinal Study of Aging (BLSA) at Johns Hopkins University and Banner Sun Health Research Institute (Banner) from the Accelerating Medicines Partnership – Alzheimer Disease (AMP-AD) consortium (https://adknowledgeportal.synapse.org/) [126, 127]. The BLSA samples consisted of 97 samples from the dorsolateral prefrontal cortex (BA9 area) representing 15 controls, 15 AsymAD and 20 AD cases [128]. The Banner samples were from prefrontal cortex of 30 controls, 28 mild cognitive impairment, 33 AsymAD and 98 confirmed AD cases [91]. The label free quantitation intensities were log2 transformed and assessed for effects from biological covariates (diagnosis, age, gender) and technical variables (batch, brain bank). We used a linear regression model accounting for biological and technical covariates. The final model used was implemented in R version 3.6.1 (R Core Team 2019) as follows:

lm(expression ∼ diagnosis + age + gender + batch + brain.bank.batch)

We evaluated the proteins that were up-regulated proteins in apoE4 lysosomes relative to apoE3 or apoE-deficient lysosomes (apoeE4>apoE3/apoE-deficient). We also evaluated the up-/down-regulated proteins in apoE4 lysosomes only relative to apoE3 lysosomes (apoE4>apoE3|apoE4<apoE3). For these protein sets, we considered their eigenprotein as the first principal component of their protein expression [129, 130]. We determined if the eigenprotein for apoeE4>apoE3/apoE-deficient or apoeE4>apoE3 were significantly different from disease diagnosis (AD, mild cognitive impairment, AsymAD) versus control using the Wilcoxon rank sum test [131]. Eigenproteins were correlated with apoE4 dosage using Pearson correlation [132].

### Statistical analysis

Statistical tests performed are described in the figure legends of their associated graphs. Generally speaking, all data presented was collected from at least three separate technical and biological replicates. In the case of iPSC-derived neurons, one differentiation batch was used as a single biological replicate. The following statistical tests were used, unless otherwise indicated: For comparisons between two conditions (one independent variable), a two-way t-test was performed. For comparisons between multiple conditions across one independent variables, a one-way ANOVA was performed followed *post-hoc* by Bonferroni’s multiple comparison test. For comparisons between multiple conditions across multiple independent variables, a two-way ANOVA was performed followed *post-hoc* by Bonferroni’s multiple comparison test. All bar graphs present data as the mean ± SEM.

## Abbreviations

Aβ: β-Amyloid
AD: Alzheimer Disease
APMS: Affinity purification mass spectrometry
apoE: Apolipoprotein E
App: Amyloid precursor protein
BafA1: Bafilomycin A1
CMA: Chaperone-mediated autophagy
COP: Coatomer
ER: Endoplasmic reticulum
ERAD: Endoplasmic reticulum-associated degradation
ERGIC: Endoplasmic reticulum-Golgi intermediate compartment
FCCP: Carbonyl cyanide p-trifluoro methoxyphenylhydrazone
GABA: Gamma-aminobutyric acid
GFP: Green fluorescent protein
GPI: Glycerophosphoinositol
GPN: Glycyl-L-phenylalanine 2-naphthylamide
HA: Hemagglutinin protein tag
iPSC: Induced pluripotent stem cells
LAMP1: Lysosome-associated membrane protein 1
Lgals3bp: Galectin 3-binding protein
LysoIP: Lysosomal immunoprecipitation
Mapt: Microtubule-associated protein tau
MR-CtsB: Magic red cathepsin B
MS: Mass spectrometry
NFT: Neurofibrillary tangles
PCA: Principal component analysis
PLA: Proximity ligation assay
PSEN1: Presenilin 1
Psen2: Presenilin 2
pTau: Phospho-tau
ROS: Reactive oxygen species
RT-qPCR: Reverse transcriptase quantitative polymerase chain reaction
SEM: Standard error of the mean
siRNA: Short interfering RNA
SRCR: Scavenger receptor cysteine-rich domain
Tau: Microtubule-associated protein tau
Tmed: Transmembrane emp24 domain-containing protein
TMEM192: Transmembrane protein 192

## Data and materials availability

All unique and stable reagents, including plasmids and LysoIP cell lines generated in this study, are available from the lead contact without restriction. The proteomic datasets supporting the conclusions of this article is available in the ProteomeXchange Consortium (http://proteomecentral.proteomexchange.org) via the PRIDE partner repository with dataset identifier PXD044942, and can be accessed using the following credentials: Username, Password: XI5oUUKC. The proteomic results and analysis sheets for both LysoIP and APMS experiments are also listed in supplemental tables S1 and S2, respectively. The human brain results published here are in whole or in part based on data obtained from the AMP-AD Knowledge Portal (https://adknowledgeportal.synapse.org/).

## Acknowledgments

We thank Robert W. Mahley for helpful discussions. We thank Ken Nakamura and Huihui Li for sharing the mtKeima plasmid, and Monther Abu-Remaileh and Ali Ghoochani for sharing LysoIP plasmids. We also thank Margaret Soucheray and Holli Deval for lab organization, Ben Polacco for bioinformatic support, and the rest of the Krogan laboratory for helpful discussions around the data presented herein. We thank the Gladstone Histology & Light Microscopy Core for supporting all confocal microscopy experiments. Human tissue samples were obtained with the help of Lea Grinberg, Wing Hung Lee, and Tia LaMore from the Neurodegenerative Disease Brain Bank, UCSF, supported by grant number P30 AG062422 (Alzheimer’s Disease Research Center). BLSA data were generated from postmortem brain tissue collected through the National Institute on Aging’s Baltimore Longitudinal Study of Aging and provided by Dr. Levey from Emory University. Banner data were provided by Dr. Levey from Emory University. A portion of these data were generated from samples collected through the Sun Health Research Institute Brain and Body Donation Program of Sun City, Arizona.

## Funding

National Institutes of Health grant R01AG059751 (NJK, DLS)

National Institutes of Health grant U54NS123746 (DLS)

National Institutes of Health grant R01AG071697 (YH)

National Institute of Neurological Disorders and Stroke grant R25NS065723 (TSC)

National Institute of Neurological Disorders and Stroke grant U24 NS072026 (Brain and Body Donation Program)

National Institute on Aging grant P30 AG19610 (Arizona Alzheimer’s Disease Core Center)

Arizona Department of Health Services contract 211002 (Arizona Alzheimer’s Research Center)

Arizona Biomedical Research Commission contracts 4001, 0011, 05-901, 1001 (Arizona Parkinson’s Disease Consortium)

Michael J. Fox Foundation for Parkinson’s Research (TSC)

California Institute of Regenerative Medicine Fellowship (EKK)

Gladstone Institutes PUMAS Program (AM)

## Author contributions

**Conceptualization:** EKK, ZC, DLS. **Data curation:** JM, LV-B, TSC. **Formal analysis:** EKK, JM, AM, AW, LV-B, TSC. **Funding acquisition:** MK, YH, NJK, DLS, TSC. **Investigation:** EKK, JM, LV-B, AM, AW, ES, GMJ, AR, EL, AA, AZ. **Methodology:** EKK, JM, GMJ, AR, EL, AZ, TSC, YH, DLS. **Project administration:** DLS. **Resources:** MK, YH, NJK, DLS. **Software:** JM. **Supervision**: EKK, TSC, MK, YH, NJK, DLS. **Validation:** EKK, JM. **Visualization:** EKK, JM, LV-B, TSC. **Writing – original draft:** EKK, DLS. **Writing – review & editing:** EKK, JM, LV-B, GMJ, TSC, YH, NJK, DLS.

## Supplemental Figures

**Figure S1:**
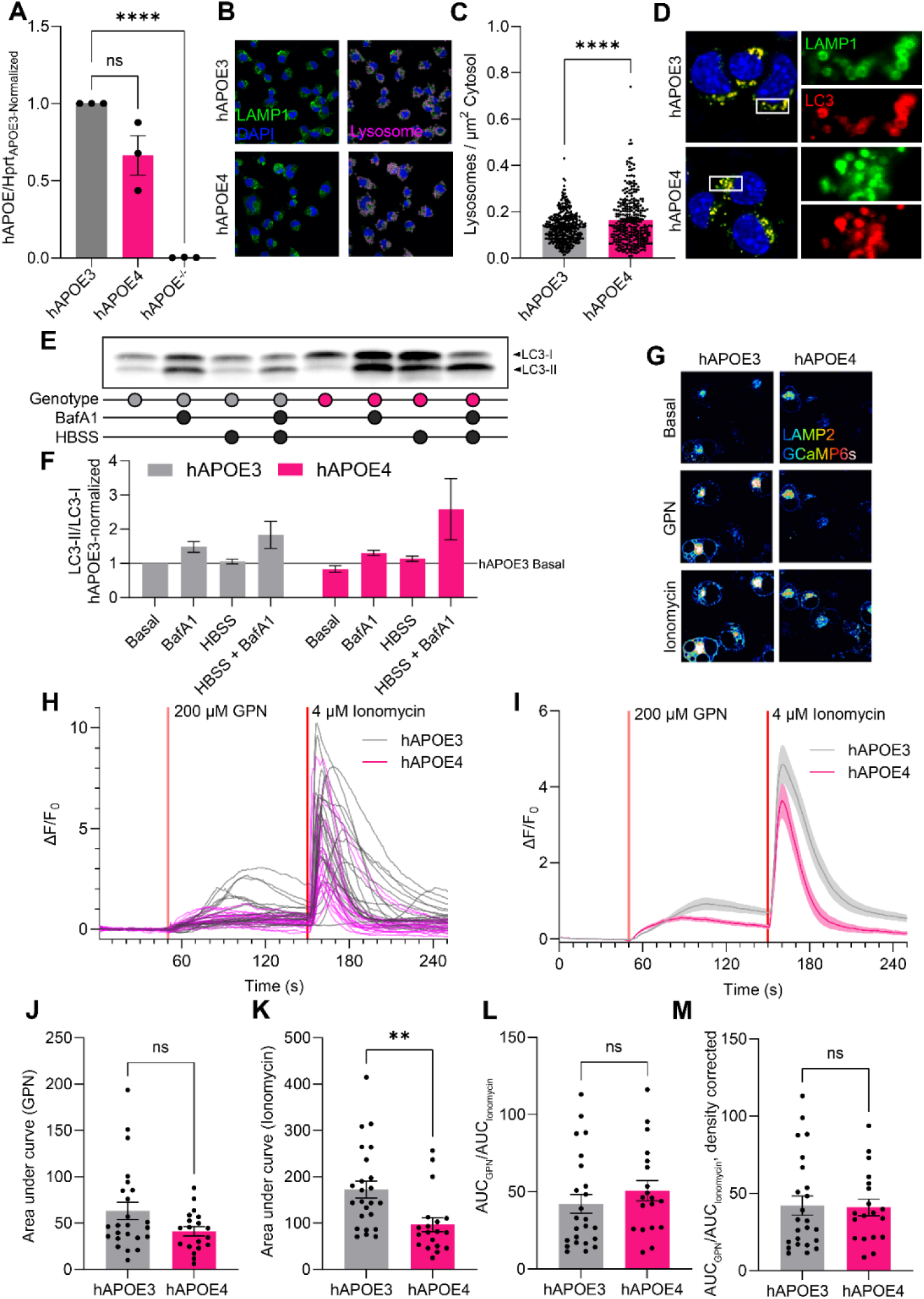
Lysosomal calcium content. (**A**) *hAPOE* gene expression across Neuro-2a cell lines, measured by RT-qPCR. (**B**) Immunofluorescence of Neuro-2a cells expressing hAPOE3 or hAPOE4, stained for the integral lysosomal membrane protein LAMP1. Right panels show puncta identification using Fiji. (**C**) Lysosomal density per cell based on images such as those in panel B. (**D**) Immunofluorescence of LAMP1 and LC3 shows lysosome-autophagosome fusion in both hAPOE3 and hAPOE4 Neuro-2a cells following 3 hours HBSS starvation and BafA1 treatment. (**E**) Representative western blot of LC3-I and LC3-II in hAPOE3 and hAPOE4 Neuro-2a cells. (**F**) Quantification of LC3-II/LC3-I ratio, normalized to internal hAPOE3 controls, across three independent replicates. (**G**) LAMP2-GCaMP6s lysosomal calcium measurements in Neuro-2a cells shown at basal states, and upon sequential stimulation using 200 µM GPN to release lysosomal calcium, and 4 µM Ionomycin to release total calcium. (**H**) Single-cell traces for lysosomal calcium release, as shown in panel D. (**I**) Mean traces of lysosomal calcium release, alongside SEM error bar traces. (**J**) Lysosomal calcium signals, as measured by the area under the curve following GPN and Ionomycin stimulation. (**K**) Total calcium sensitivity of apoE3 and apoE4 cells, as measured by the area under the curve following Ionomycin stimulation. (**L**) Relative lysosomal calcium release, normalized to the calcium sensitivity. (**M**) Relative lysosomal calcium release normalized to the lysosomal density, as calculated in panel C, estimates the lysosomal calcium content of each individual lysosome.

**Figure S2:**
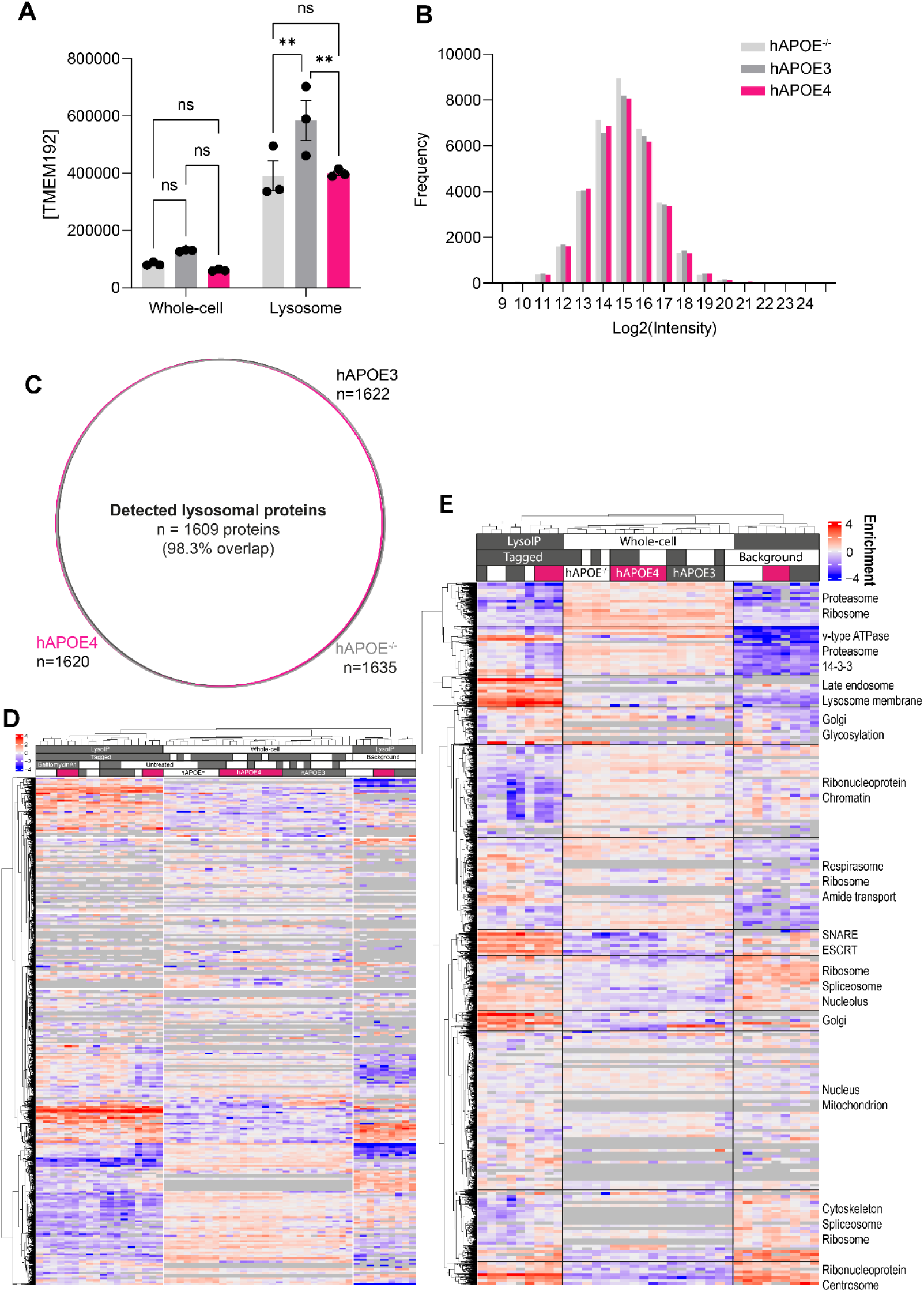
Abundance proteome coverage and clustering across apoE genotypes. (**A**) Human lysosomal TMEM192 tag intensities in whole-cell and lysosomal fractions of untreated Neuro-2a cells from 3 independent experiments. (**B**) Histogram of Log2-transformed protein group intensities across all obtained Neuro-2a lysosomal proteomes, stratified by genotype. (**C**) Venn diagram of lysosome-enriched proteins detected in LysoIP samples across genotypes. (**D**) Full heat-map generated from raw intensities of all detected proteins across all samples reveals distinct clustering of LysoIP, whole-cell, and LysoIP background datasets, as well as sub-clustering based on Bafilomycin treatment and apoE genotype. (**E**) Heat map excluding BafA1-treated samples, including cellular component annotations of detected proteins. Groups of data were analyzed in GraphPad Prism by 2-way ANOVA followed *post-hoc* by Dunnett’s multiple comparisons test. **P<0.01.

**Figure S3:**
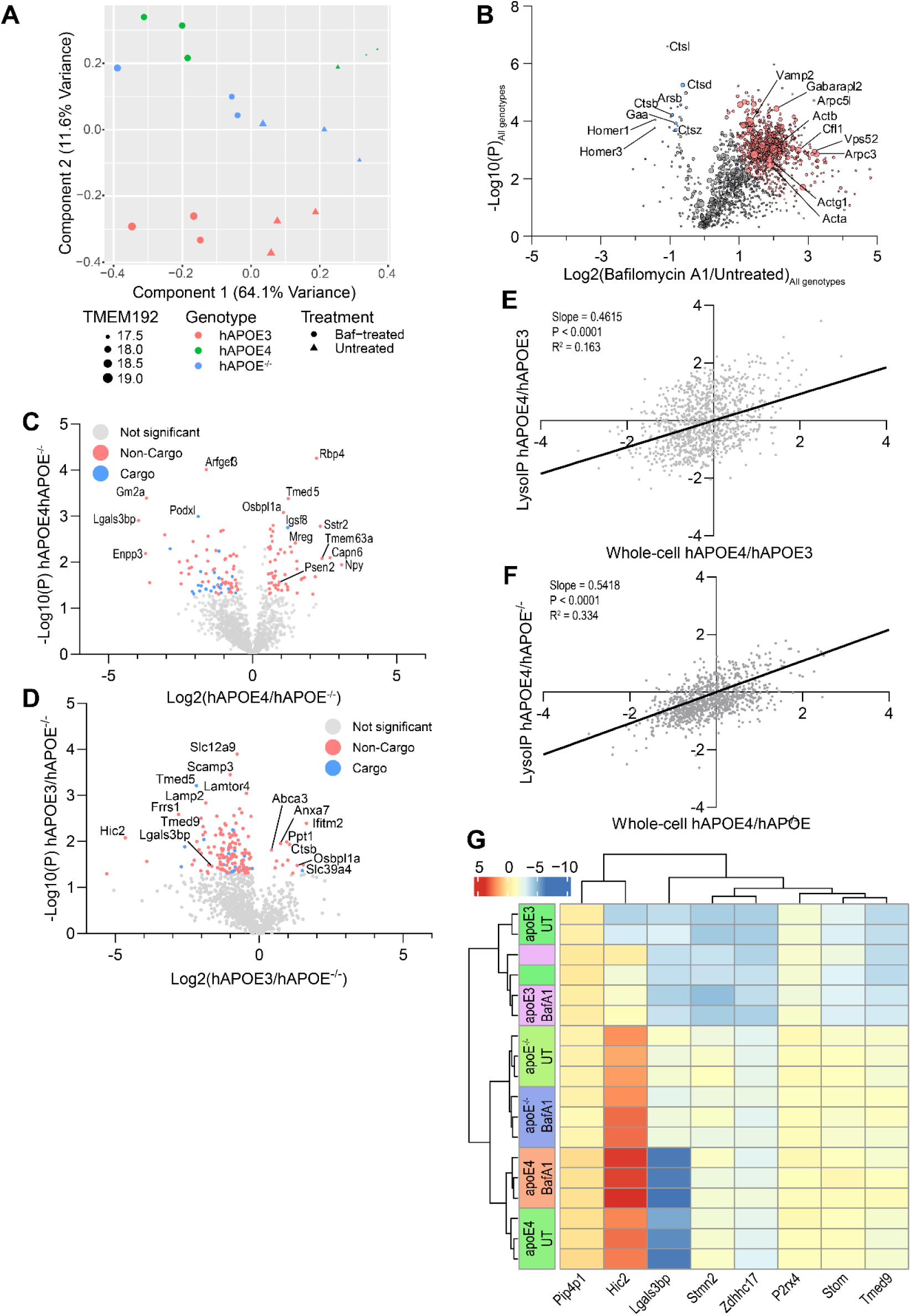
Proteomic comparisons of apoE lysosomes. (**A**) Principal component analysis of all lysosomal proteomes separates samples based on Bafilomycin treatment (component 1) and apoE genotype (component 2). (**B**) Volcano plot comparing BafA1 and untreated lysosomal proteomes, pooled across hAPOE genotypes. Node sizes indicate relative protein abundances following BafA1 treatment. (**C-D**) Volcano plot comparing hAPOE4 and hAPOE^−/−^(C) or hAPOE3 and hAPOE^−/−^ (D) lysosomal proteomes. Proteins significantly changed in hAPOE4 lysosomes are colored by their status as a lysosomal cargo protein, based on basal (red) or BafA1-dependent (blue) lysosomal enrichment. (**E-F**) Comparison of hAPOE4-associated changes to whole-cell and lysosomal protein abundances to hAPOE3 (E) and hAPOE^−/−^ (F) proteomes, showing positive correlation between changed whole-cell and lysosomal protein levels. Only proteins detected across four sample sets (genotype and fraction) were be plotted. (**G**) Partial least squares-discriminant analysis of lysosomal proteomes reveals proteins separating experimental groups. In particular, hAPOE4 was accompanied by lysosomal Lgals3bp depletion.

**Figure S4:**
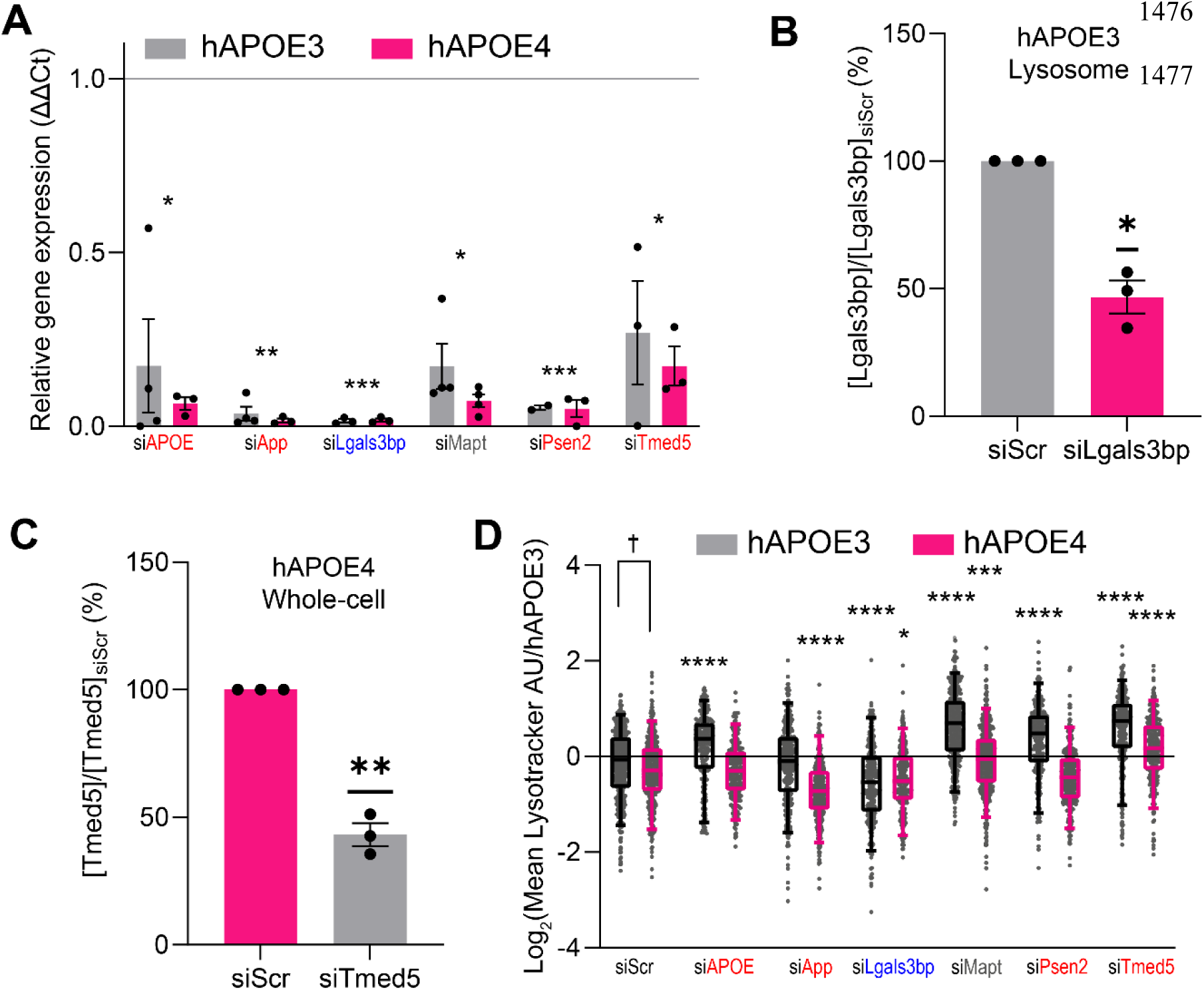
Identification of proteomic drivers underlying hAPOE4 lysosomal defects. (**A**) Gene expression upon targeting indicated genes with siRNAs in Neuro-2a cells. Expression level changes are indicated relative to *Hprt* expression and siScr-treated cells. (**B**) Efficiency of Lgals3bp protein reduction in hAPOE3 lysosomes upon Lgals3bp siRNA knockdown relative to scrambled siRNA controls as measured by mass spectrometry. Lysosomal Lgals3bp was measured since it was undetected in the whole-cell lysate upon knockdown, precluding calculation of its knockdown efficiency. (**C**) Efficiency of Tmed5 protein reduction in hAPOE4 whole-cell lysate upon Tmed5 siRNA knockdown relative to scrambled siRNA controls as measured by mass spectrometry. (**D**) Influence of gene silencing on LysoTracker staining, normalized to hAPOE3 LysoTracker intensity. Knockdown comparisons are performed relative to the same genotype siScr control. Dagger (†) comparison shows difference between hAPOE3 and hAPOE4 cells. Data points indicate individual cells from three independent experiments. Groups of data were analyzed in GraphPad Prism by 2-way ANOVA followed *post-hoc* by Dunnett’s multiple comparisons test. Pairs of data were analyzed by t-tests. *P<0.05, **P<0.01, ***P<0.001, ****P<0.0001.

**Figure S5:**
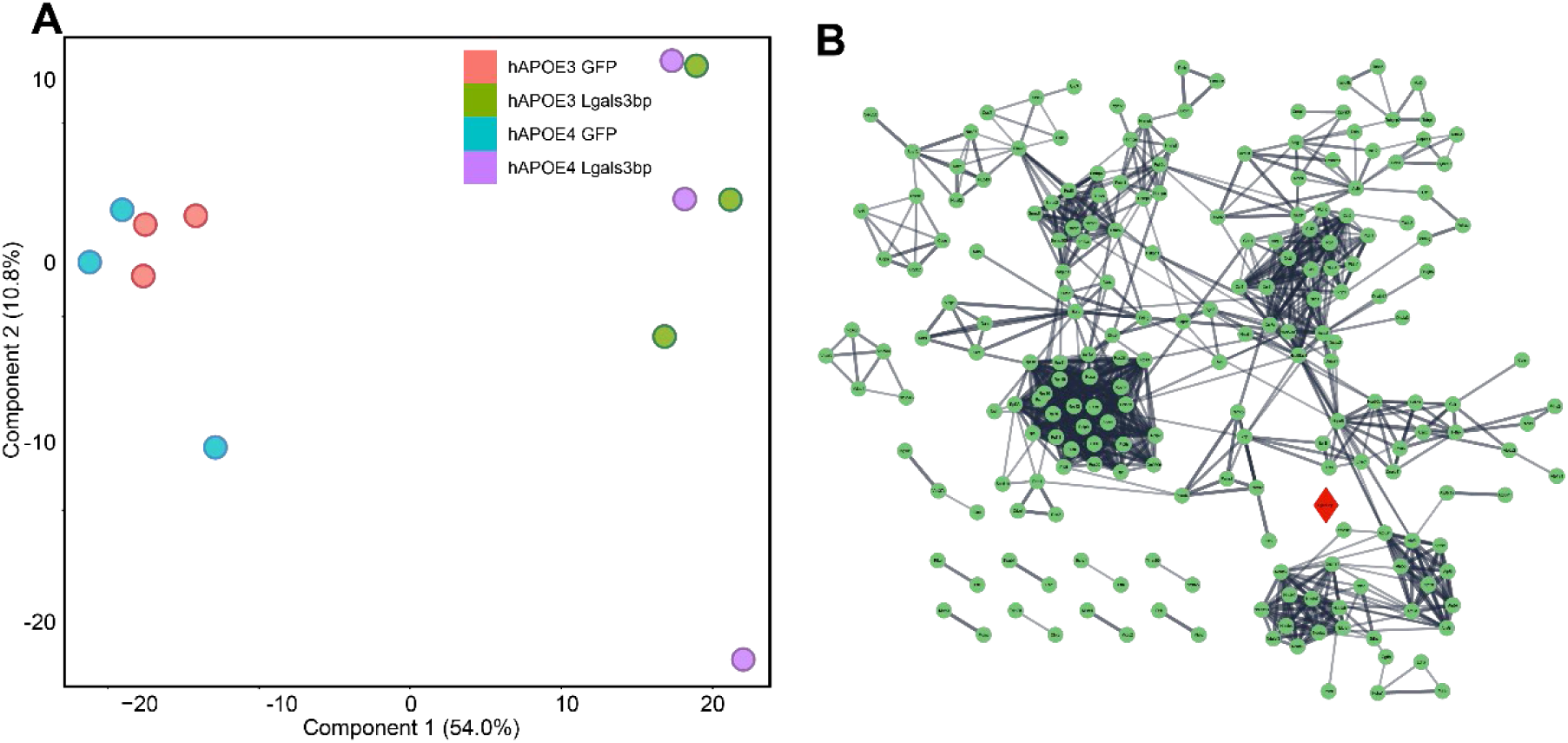
The neuronal Lgals3bp interactome. (**A**) Principal component analysis of affinity-purified proteomes shows clustering of Lgals3bp-GFP-purified samples distinct from GFP background samples (component 1), and slight separation of apoE3 and apoE4 samples (component 2). (**B**) Physical STRING subnetwork of identified Lgals3bp interactors. Lgals3bp is highlighted in red.

**Figure S6:**
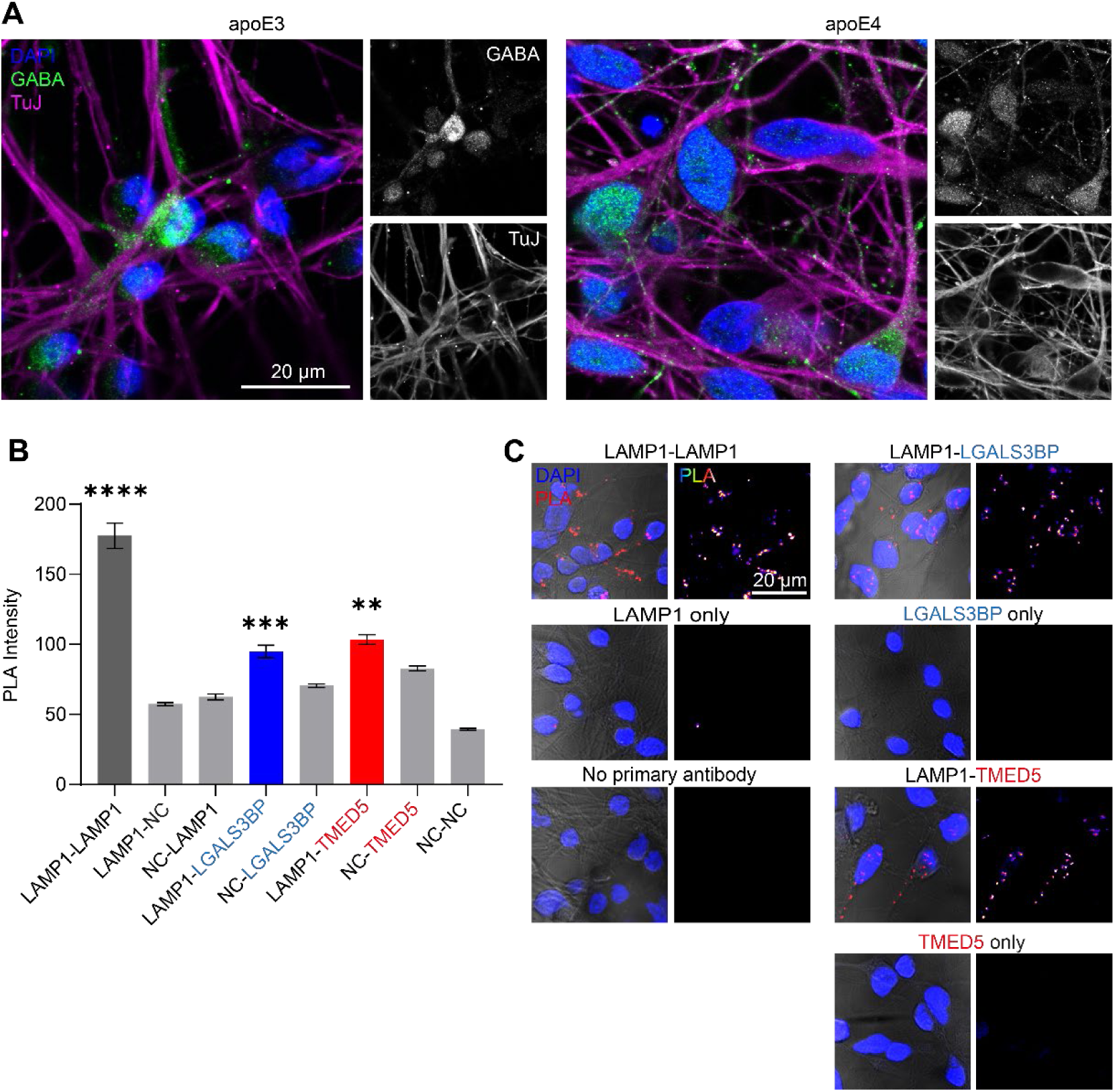
Controls for iPSC-derived neuron proximity ligation assay. (**A**) Immunofluorescen1c4e8o3f neuronal markers following 49 days of differentiation, revealing pure cultures of TuJ^+^ neurons including GABA^+^ interneurons for both genotypes. (**B**) Establishing PLA sensitivities, showing significant signals above single antibody negative controls (NC) for LAMP1-LAMP1, LAMP1-LGALS3BP, and LAMP1-TMED5 assays. (**C**) Representative images for quantification shown in panel B. Individual cells from three independent experiments were analyzed by a one-way ANOVA followed by a *post-hoc* Bonferroni multiple comparison test between dual antibody samples and single-antibody controls; the least statistically significant comparison is highlighted. **P<0.01, ***P<0.001, ****P<0.0001.

